# Variation in host home range decreases rabies vaccination effectiveness by increasing the spatial spread of rabies

**DOI:** 10.1101/815100

**Authors:** Katherine M. McClure, Amy T. Gilbert, Richard B. Chipman, Erin Rees, Kim M. Pepin

**Affiliations:** United States Department of Agriculture, Animal and Plant Health Inspection Service, Wildlife Services, National Wildlife Research Center, Fort Collins, Colorado, USA; Colorado State University, Department of Microbiology, Immunology, and Pathology, Fort Collins, Colorado, USA; United States Department of Agriculture, Animal and Plant Health Inspection Service, Wildlife Services, National Rabies Management Program, Concord, New Hampshire, USA; Land and Sea Systems Analysis Inc., Granby, Québec, Canada; Public Health Risk Sciences Division, National Microbiology Laboratory, Public Health Agency of Canada, Saint-Hyacinthe, Québec, Canada

**Keywords:** movement, ORV, rabies, raccoon ecology, spatially-explicit model, vaccination

## Abstract

1. Animal movement influences the spatial spread of wildlife infectious diseases through host-host contact structure and hence pathogen transmission. Wildlife disease hosts vary in characteristics related to pathogen transmission, which can increase the spread and intensity of disease outbreaks. The consequences of home range size variation on wildlife disease dynamics are poorly understood, but could help to predict disease spread and determine more effective disease management strategies.
2. We developed a spatially-explicit individual-based model to examine the effect of variation in host home range size on the spatial spread rate, persistence, and incidence of rabies virus (RABV) in raccoons (*Procyon lotor*). We tested the hypothesis that host home range area variation decreases vaccination effectiveness in wildlife host populations following pathogen invasion into a vaccination zone.
3. We simulated raccoon demography and RABV dynamics across a range of magnitudes and variances in weekly home range radius distributions for raccoons, and compared results to conditions that assumed a fixed host home range area. We examined how variable host home range radius distributions influenced the relative effectiveness of three components of orally-baited raccoon RABV vaccination (ORV) programs—timing and frequency of bait delivery, width of the zone where ORV baits were delivered, and proportion of hosts immunized.
4. Variability in home range radius distributions increased RABV spread rates by 1.2 - 5.2-fold compared to simulations with fixed radii. More variable host home range radius distributions decreased relative vaccination effectiveness by 71% compared to less variable host home range radius distributions under conventional vaccination conditions. We found that vaccination timing was more influential for vaccination effectiveness than vaccination frequency or vaccination zone width.
5. Our results suggest that variation in wildlife home range exploration increases the spatial spread and incidence of wildlife disease. Our vaccination results underscore the importance of prioritizing individual-level space use data collection to understand the dynamics of wildlife diseases and plan their effective control and elimination.

## INTRODUCTION

Animal movement is a key component of many ecological processes, including population dynamics, species interactions, and the spatial spread of wildlife infectious diseases (Bowler & Benton, 2005; Hess, 1996; Kays, Crofoot, Jetz, & Wikelski, 2015). Natural and human-mediated movements of infected domestic animals and wildlife have been implicated in the spread of diseases such as bovine tuberculosis (TB) in cattle and possums, chronic wasting disease in mule deer, and rabies in raccoons (Corner, Stevenson, & Collins, 2003; Farnsworth, Hoeting, Hobbs, & Miller, 2006; Gilbert et al., 2005; Rosatte et al., 2006). In diseases with direct transmission, host movement influences the spatiotemporal distribution of host-host contact, and underpins the contact structure between infectious and susceptible individuals (Morales et al., 2010). Animal movement can play critical direct and indirect roles in pathogen transmission, yet our understanding of how spatiotemporal or individual-level differences in natural wildlife host movement affects disease dynamics is limited.

Effects of variation in wildlife movement on the transmission of directly-transmitted wildlife pathogens depend on the interplay of host ecology, pathogen ecology, and the spatial structure of host contact. Host variability in contact rates, susceptibility, infectiousness, or spatiotemporal variability in other host characteristics related to pathogen transmission can increase both the intensity of disease outbreaks and probability of pathogen extinction (Lloyd-Smith, Schreiber, Kopp, & Getz, 2005; Woolhouse et al., 1997), and are common in both humans and wildlife (Paull *et al*. 2012; VanderWaal and Ezenwa 2016). Contact heterogeneities in wildlife can arise from complex social structure or fluctuations in spatial host distributions (Craft, 2015; Drewe, 2010; Hamede, Bashford, McCallum, & Jones, 2009), and consistent individual-level variation in wildlife movement related to natal dispersal and foraging tactics have recently been documented (Bonnot et al., 2015; Plowright et al., 2017). Simulations of personality-dependent individual-level movement variation in animals suggest such variation shape animal contact rates (Spiegel, Leu, Bull, & Sih, 2017). If spatiotemporal variation in host movement promotes heterogeneity in the capacity for individuals to contact or transmit pathogens to other hosts, host movement variation could result in transmission heterogeneity that increases wildlife disease spread and incidence while decreasing pathogen persistence. Understanding the effects of host movement variation on wildlife disease dynamics could thus be critical for predicting spatial spread (Cross et al., 2010).

Host movement also influences the effectiveness of wildlife disease intervention strategies. For example, the explicit consideration of red fox (*Vulpes vulpes*) movement and territoriality in oral-baited vaccination efforts was crucial for the elimination of red fox rabies in Western Europe, reinforcing the importance of understanding host ecology when targeting free-ranging wildlife species for disease elimination (Freuling et al., 2013; Murray et al., 1986). Conversely, disease management strategies can affect animal movement and lead to unintended consequences for pathogen transmission. For instance, badger culling to reduce spillover of bovine TB to cattle in the UK increased badger movement, contact rates, and transmission to cattle near the culling zone (Donnelly et al., 2006; Pope et al., 2007). Ultimately, targeted control measures that treat or eliminate individuals that are most connected through movement could be more effective than applying interventions randomly (Lloyd-Smith et al., 2005). In this context, an understanding of how variation in and scope of host movement influence wildlife disease management strategies appears to be important for planning effective vaccination efforts.

Rabies (RABV) is a zoonotic infectious disease caused by ssRNA viruses of the genus *Lyssavirus* (Wunner, 2007). Transmission occurs primarily through direct contact among hosts, and infected mammals almost invariably develop fatal encephalomyelitis (Rupprecht, Hanlon, & Hemachudha, 2002). Raccoon RABV is the most prevalent terrestrial RABV variant in the U.S., with raccoons (*Procyon lotor*) accounting for the highest proportion of reported RABV cases in terrestrial wildlife species from 1991-2014 (Ma, 2018). One primary objective of U.S. RABV management efforts currently focuses on preventing the westward expansion of raccoon RABV from the eastern U.S., primarily by deploying oral RABV vaccine baits containing live recombinant vaccines that provide long-term immunity for raccoons to RABV infection when ingested (Blanton et al., 2018; Slate et al., 2009). Oral RABV vaccination (ORV) has proven effective for the elimination of canine RABV in coyotes in the U.S., raccoon RABV in Canada, and red fox RABV throughout Western and Central Europe (Müller et al., 2015; Rosatte et al., 2009; Sidwa et al., 2005).

We developed a spatially-explicit individual-based model (IBM) of raccoon population dynamics and raccoon RABV transmission to investigate how spatiotemporal variation in wildlife host movement (implemented as weekly home range size variation) affects the spatial spread, persistence, and incidence of wildlife disease and vaccination effectiveness. We hypothesized that variable home range size would increase pathogen spread and incidence rates, and decrease pathogen persistence, compared to conditions assuming constant patterns of host home range size, as would be predicted by super-spreader theory (Lloyd-Smith et al. 2005). We tested the hypothesis that variation in host home range size decreases vaccination effectiveness in wildlife host populations following the invasion of RABV into an ORV zone, and examined the relative effectiveness of ORV strategies targeting raccoons. We predicted that fall vaccination would be more effective than late-spring vaccination because young of the year would be old enough to consume ORV baits in the fall but not in the late spring (Wandeler, 1991). We also predicted that the probability that RABV will breach the vaccination zone and enter the RABV-free region would decrease with increasing width of the vaccination zone.

## METHODS

### I. Model design

#### A. Approach

We used a spatially-explicit, discrete-time IBM approach to examine the role of variable home range size in spatial spread of RABV and ORV effectiveness, within the context of other realistic complexities in raccoon ecology such as social structure and natal dispersal. We compared the effects of fixed home range sizes to variable home range sizes across a range of similar magnitudes of home range size to identify when home range size variation had the strongest effects on spatial spread rates and ORV effectiveness. Variation in home range size was implemented as week-to-week stochastic changes in home range radii, compared against constant weekly home range radii. We describe components of the model below (spatial context, demographic dynamics, social structure, contact structure, disease transmission, and vaccination), and provide additional details using the updated Overview, Design Concepts, and Details protocol for IBMs (Grimm et al., 2010) in the Supplementary Information.

#### B. Spatial context

Simulated landscapes contained 1km^2^ gridded cells comprising four spatially consecutive zones: seeding (1 × 20 km), spreading (10 × 20 km), vaccination (20-60 × 20 km), and breach (10 × 20 km), for a total landscape area ranging between 820 km^2^ and 1620 km^2^ (Fig. 1A). The cell-level was discrete with a carrying capacity of 15 individuals/km^2^—corresponding to typical suburban raccoon densities (Table S1). Individuals were assigned a randomly drawn home range centroid point (i.e., in continuous space) that occurred within a disrcete grid cell. We tracked disease-related and demographic characteristics of each individual on a weekly time-step. Simulations were conducted in Matlab R2016b (Version 9.1.0, MathWorks, Inc., Natick, MA). Results were analyzed in R v3.4.2 (R Core, 2017).

**Figure 1.**
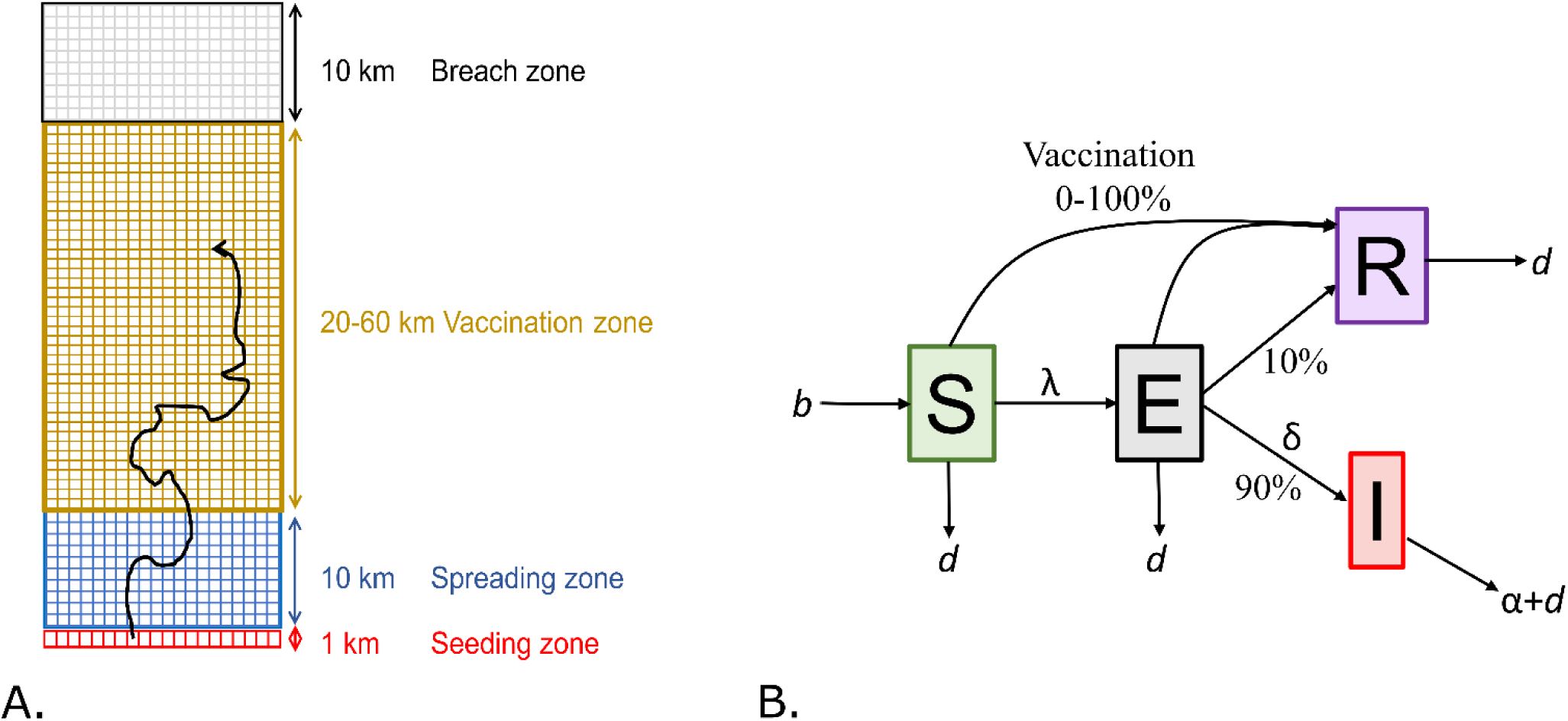
Landscape and disease components of model. **A)** The landscape consisted of 1km^2^ grid cells with 4 zones. RABV was introduced to the seeding zone; breach occurred when an infectious individual crossed into the breach zone. **B)** Transitions between S (susceptible), E (exposed), I (infectious), and R (recovered) disease states are governed by the force of infection (*λ*), incubation rate (δ), and disease-induced mortality rate (α). Demographic rates include birth (*b*) and natural death (*d*).

#### C. Population turnover, dispersal, and social structure

We modeled reproduction as a single six-week birth pulse from April to mid-May (Table S1 and SI1 Methods for more details). Individuals were subject to density-dependent mortality to maintain densities at or below within-grid carrying capacity. If grid-level carrying capacity was exceeded, individuals within the grid were randomly chosen and removed from the simulation, with younger individuals taken first to mimic observed patterns in juvenile and adult survivorship (Gehrt & Fritzell, 1999). We modeled male-biased natal dispersal as two random variables, natal dispersal distance and dispersal age (described in SI1). Individuals that moved off the landscape during dispersal were lost permanently, and did not move back onto the landscape. We modeled social structure comprising family groups of females and offspring, male dyads, and solitary males. Field and genetic studies suggest that daughters may associate with mothers and her offspring into adulthood (Cullingham et al., 2008; S. Gehrt & Fritzell, 1998), while males often associate in relatively long-lasting, non-familial dyads followed by separation to become independent as they mature (Gehrt, Gergits, & Fritzell, 2008; Gehrt & Fritzell, 1998b). We assumed that individuals in the same family group had the same home range centroids (Table S1).

#### D. Weekly contact

We modeled variation in home range size as a random variable described by a gamma distribution of weekly-varying home range radii distances using parameters derived from (or similar to) weekly variation in GPS collar data (in km, described below, Table S1). This radius was randomly assigned to each susceptible individual relative to their home range centroid at each time step, and delineated a circular home range area (Fig. S1). We assumed individuals explored the entirety of their weekly home range areas. For scenarios with constant home range radii, all individuals were assigned the same home range radius that was explored entirely weekly. Contact opportunities in the home range were assumed equally probable given the high degree of social connectivity observed in suburban raccoon populations (Hirsch, Prange, Hauver, & Gehrt, 2013).

#### E. Disease transmission

We modeled disease dynamics of rabies with four disease states: susceptible (S), exposed but not infectious (E), infectious (I), and recovered (R) (Fig. 1B). Density-dependent transmission occurred when home range centroids of infectious individuals were within the weekly home range area of a susceptible individual, according to a fixed transmission probability given contact. Transmission probability was based on family group membership to reflect potential differences in within- vs. between-group contact rates in raccoons (Fig. S1). Within-group transmission probability was fixed at 0.5, while between-family transmission probability ranged between 0.001 - 0.5. Within-group transmission was non-spatial because we assumed weekly contact probability was 100% within family groups, whereas between-group transmission was spatially-explicit because contact required that home range centroids of infectious individuals were in the weekly home range of a susceptible individual. Recovery rate of exposed individuals was 10% to capture variation in levels of acquired rabies immunity observed in raccoons (Slate et al., 2014). Disease-induced mortality was 100% for infectious individuals (Hanlon, Niezgoda, & Rupprecht, 2007), and occurred one week after individuals transitioned from the exposed to infectious disease class (Hanlon et al., 2007).

The force of infection, or the per capita rate at which susceptible individuals seroconvert to the exposed class, *λ_t_* at week *t*, was:

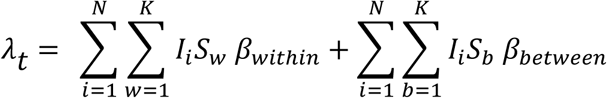

where *β_within_* is within-group transmission probability, *β_between_* is between-group transmission probability, *w* represents individuals in the same family group as *Ii* in week *t*, and *b* represents individuals that are not in the same family group as focal individual *Ii* in week *t* but are located within the weekly home range of *Ii.* RABV incubation period was drawn from a Poisson distribution (mean = 4 weeks; Table S1).

#### F. Vaccination

To model vaccination, we randomly selected a fixed proportion of animals within the vaccination zone irrespective of disease or vaccination status, and seroconverted susceptible and exposed individuals, with a two-week lag for development of vaccine-induced immunity. We neglected factors that affect achieved vaccination coverage (e.g., non-target competition for baits, bait shyness, operational limitations), and assumed that any particular coverage could be achieved, because we were interested in exploring how home range size variation affected breach probability over a wide range of theoretical coverage levels. We assumed that vaccinated individuals acquired lifetime immunity with no waning. Individuals younger than 17 weeks were not vaccinated because delivery was by oral vaccine baiting and raccoons at this age may still be dependent on the dam for nourishment (Montgomery, 1969). Vaccination was assumed ineffective on infectious individuals.

### II. Movement data and analysis

To estimate host home range movement distributions, we obtained GPS location data from 26 GPS-collared free-ranging raccoons captured in a suburban ORV area of Burlington, VT. GPS locations, or fixes, were recorded every 30 minutes to 2 hours from 6pm to 6am from late July through mid-September 2016. For each individual, we calculated the maximum distance (in km) between all fixes within a week. We fit a gamma distribution to observed distances using maximum likelihood methods (mean = 0.82 km, median = 0.75 km and variance = 0.16), considering these distances a measure of an individuals’ maximum weekly exploratory potential. We used this gamma distribution as a random variable describing the weekly-varying home range radius in our vaccination simulations. We used a second, theoretical gamma distribution with a higher variance to explore how more variable home range movement affected vaccination effectiveness (mean = 1km, median = 0.84 km, variance = 0.5).

### III. Simulations

All simulations included a one-year demographic transient period followed by the exposure of all hosts located in the middle grid cell of the seeding zone to rabies (∼15 individuals, early spring, week 11). Sensitivity analyses included a full factorial design of three parameters: 1) shape and 2) scale parameters of the weekly home range radius gamma distribution, and 3) between-group transmission probability. We used four scale parameters of the gamma distribution (0.2, 0.5, 1, and 2) corresponding to increasing variance with the associated shape parameters that correspond to medians of the gamma distribution ranging from 0.2 - 3km in 0.2 increments (Fig. S3). Between-group transmission probabilities were evaluated from 0.001 - 0.5 in logarithmic intervals for a total of 600 parameter sets. For static home range movement simulations, we examined effects of the fixed home range radius on outbreak dynamics by varying the home range radius from 0.2 – 3km in 0.2 increments for an additional 150 parameter sets. Simulations occurred on a 1220km^2^ landscape without vaccination. We ran 100 eight-year simulations per parameter set for a total of 75,000 simulations.

To explore the effect of host movement variation and ORV strategies on the probability that rabies will breach a vaccination zone, we modeled vaccination in a separate set of simulations. We ran all combinations of two between-group transmission probabilities (0.05 and 0.1), two weekly home range radius distributions (described above), and three components of ORV deployment: 1) vaccination coverage -proportion of animals immunized within the ORV zone, 2) timing and frequency of vaccine application, and 3) ORV zone width (Fig. 2). We examined three vaccination deployment strategies that have been employed previously for ORV deployment (fall, spring, or both fall and spring). We explored three vaccination zone widths (20km, 40km, and 60km, Figs. 1A & 2), and modeled vaccination coverage ranging from 0 - 100% in 10% increments, where 0% was the no vaccination control (Fig. 2). Vaccination coverage is a key component of successful vaccination efforts as it underlies herd immunity, or the population-level immunity required for a pathogen to decline (Fine, 1993). We ran 396 unique parameter sets, with 100 ten-year replicate simulations per parameter set.

**Figure 2.**
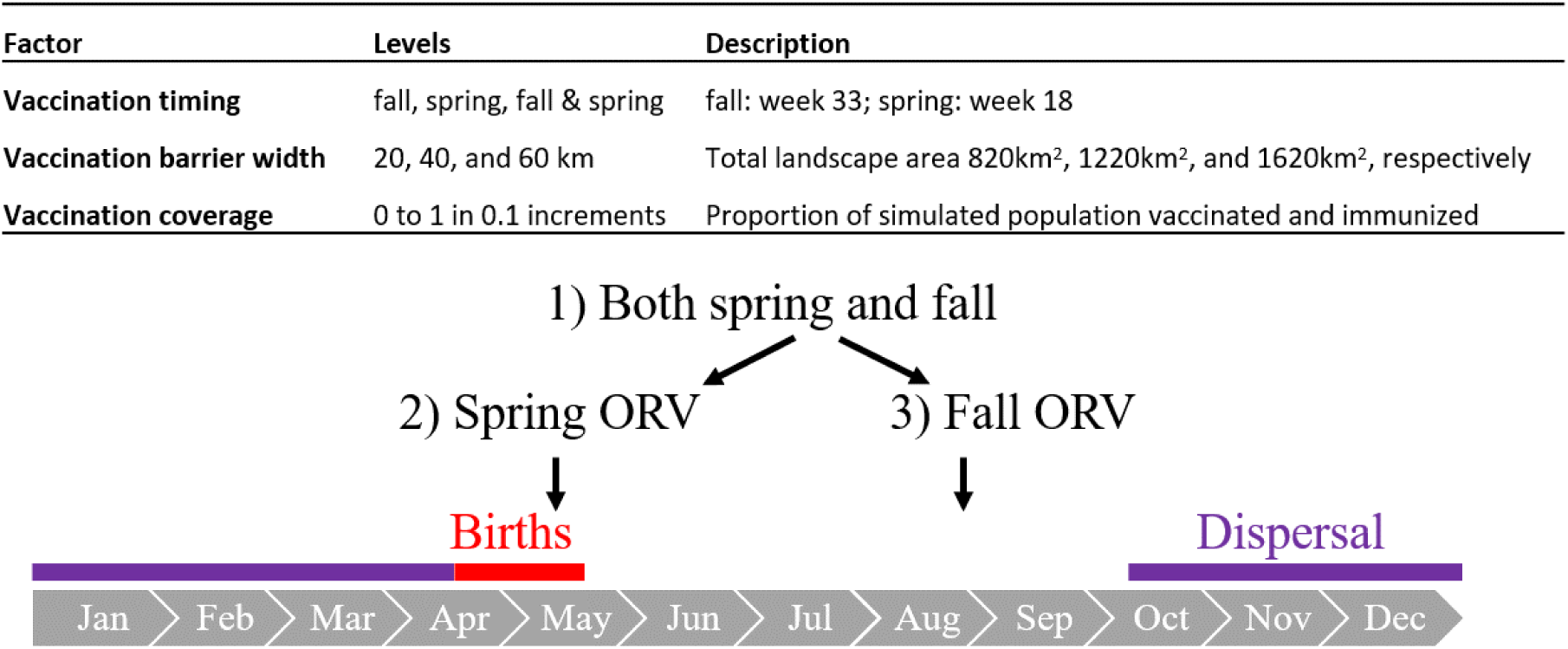
Components of vaccination. The timing and frequency of ORV (oral rabies vaccination) in relation to the annual birth pulse and male-biased natal dispersal is shown below.

### IV. Model outputs and statistical analysis

#### Sensitivity analyses

We calculated annual spatial spread rate, pathogen persistence, and per capita annual incidence as outputs of our sensitivity analyses. We calculated annual spatial spread rate (km/year) as the linear distance that RABV traveled/year from the seeding zone. We restricted spread calculations to simulations where annual incidence rate was ≥ 0.001 because we were interested in simulations that led to ongoing transmission (i.e., avoided stochastic fade-out at initiation). We defined RABV persistence as the presence of at least one exposed or infectious individual in any cell at the last time step of the eight-year simulation. Annual incidence rate was calculated as the mean annual new cases/annual maximum population size across years in which infections were present, constrained to runs where annual incidence rate was ≥ 0.001. We analyzed these outputs for fixed and variable weekly home range radii separately using generalized linear models (GLM), with home range size variation, magnitude, transmission probability, and their interactions as covariates (see Table S2 for model specification).. Specifically, covariates included median distance of the home range radius distribution (or in static home range case, the value of the constant home range radius), the scale parameter of the gamma distribution (for variable home range movement simulations only), and between-group transmission probability *Vaccination analyses*. We defined a RABV breach as a binary response variable in which infectious individuals did or did not breach the vaccination zone during the simulation (Fig. 1A, binomial GLM with logit link). We report breach probability as the proportion of 100 simulations in which the vaccination zone was breached. Fixed GLM effects included vaccination timing, coverage, zone width, between-group transmission probability, and weekly home range radius distribution.

#### Model evaluation and R_0_

For sensitivity and vaccination simulations, we evaluated the relative support of covariates using Akaike Information Criterion (AIC; Akaike, 1973), including all two-way interactions, and describe relationships of responses to covariates using the best supported model for each response variable (SI1). We calculated vaccination effectiveness as 1 – *v*, where *v* is the minimum vaccination coverage required to reduce RABV breach probability to zero. Lastly, we calculated R_0_, the average number of transmissions from an index case in a completely susceptible population, across 1,000 two-year replicate simulations using the data-informed home range radius distribution and transmission probability = 0.05 (see SI1 for further details).

## RESULTS

### Effects of home range size variation on spatial spread, persistence, and incidence

Variation in the weekly home range radius distribution increased spatial spread rates across all magnitudes of home range radii relative to simulations assuming fixed home range radii (Figs. 3A-D & 4A, Tables S2 & S3). For home range radii between 0.03–3km, the less variable distributions (scale parameter of the gamma distribution = 0.5, mean variance range = 0.33–0.88, inset Fig. 5B) increased spatial spread rates by 176%, while more variable distributions (scale parameters of the gamma distribution = 1 and 2, mean variance range = 1.91–4.41, Figs. 3C & D) increased spatial spread rates by 282-518% across all transmission probabilities. The relative effect of increased variance in host range radius distributions compared to fixed conditions was most pronounced at lower median home range radii (Fig. S4A), suggesting variable space usage may most strongly affect spread rates at smaller home range areas. However, this spatial spread pattern may also be influenced by the relative difference in variance in high and low variance models, which is inversely related to the median size of home ranges (Fig. S5). Figures 3A-D also demonstrate the effects of transmission probability on increasing rates of spatial spread, highlighting the interactive effect of home range size and transmission probability on spatial spread rates (Table S2).

**Figure 3.**
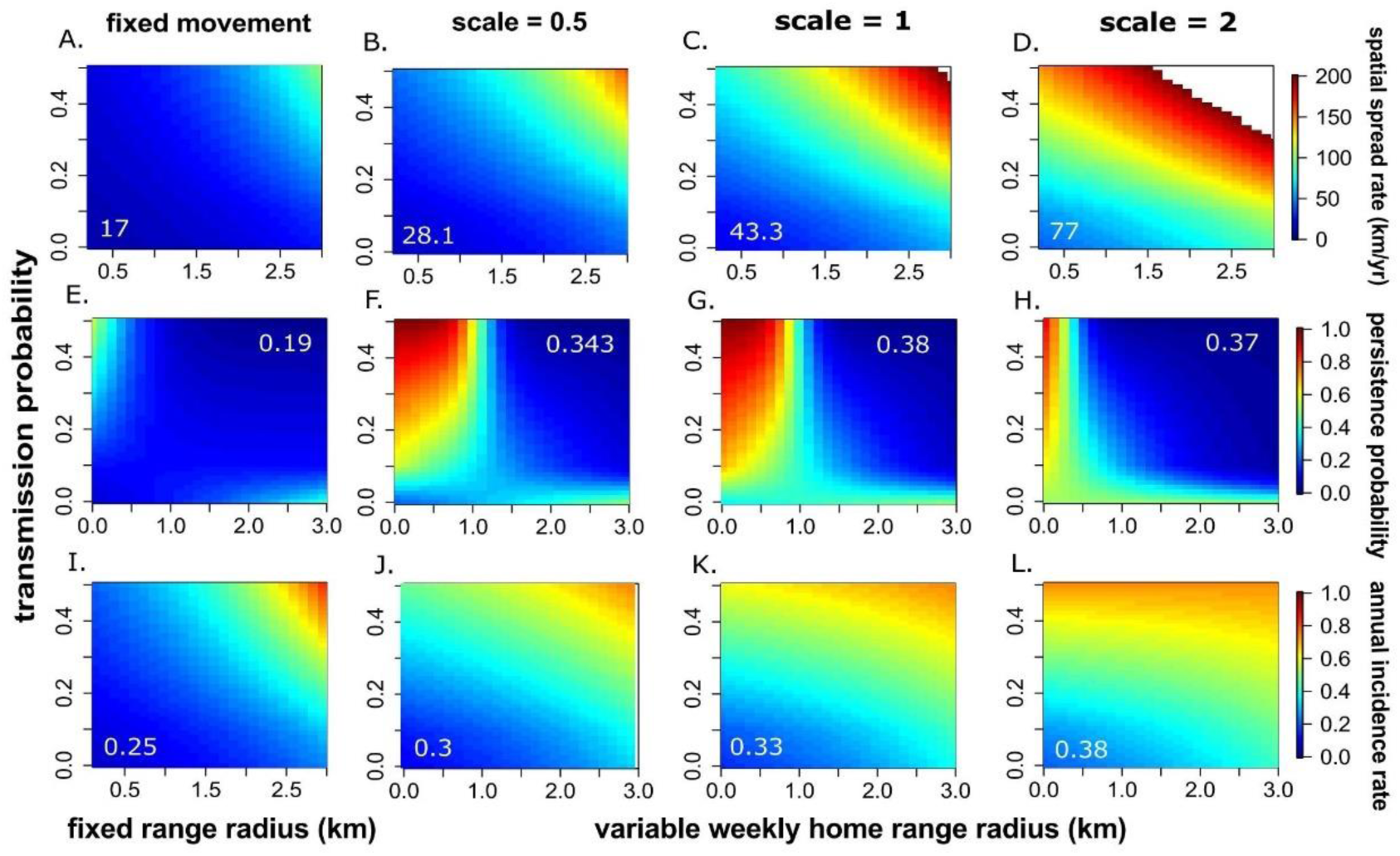
Annual spatial spread, persistence probability, and annual incidence rate. Rows correspond to model outcomes with legends shown on the right. Columns correspond to different levels of variation in home range area, beginning with fixed home range radius distance (km) on the left followed by increasing values of the scale parameter of the host home range radius distribution, reflecting increasing variance in host home range movement. Each plot shows the same range of median home range radius along the X-axis. Heat map colors are: **A-D)** annual spatial spread rate (km/year), **E-H)** pathogen persistence probability over the eight-year simulations, and **I-L)** mean annual incidence rate. The mean value across all simulations is shown in white.

**Figure 5.**
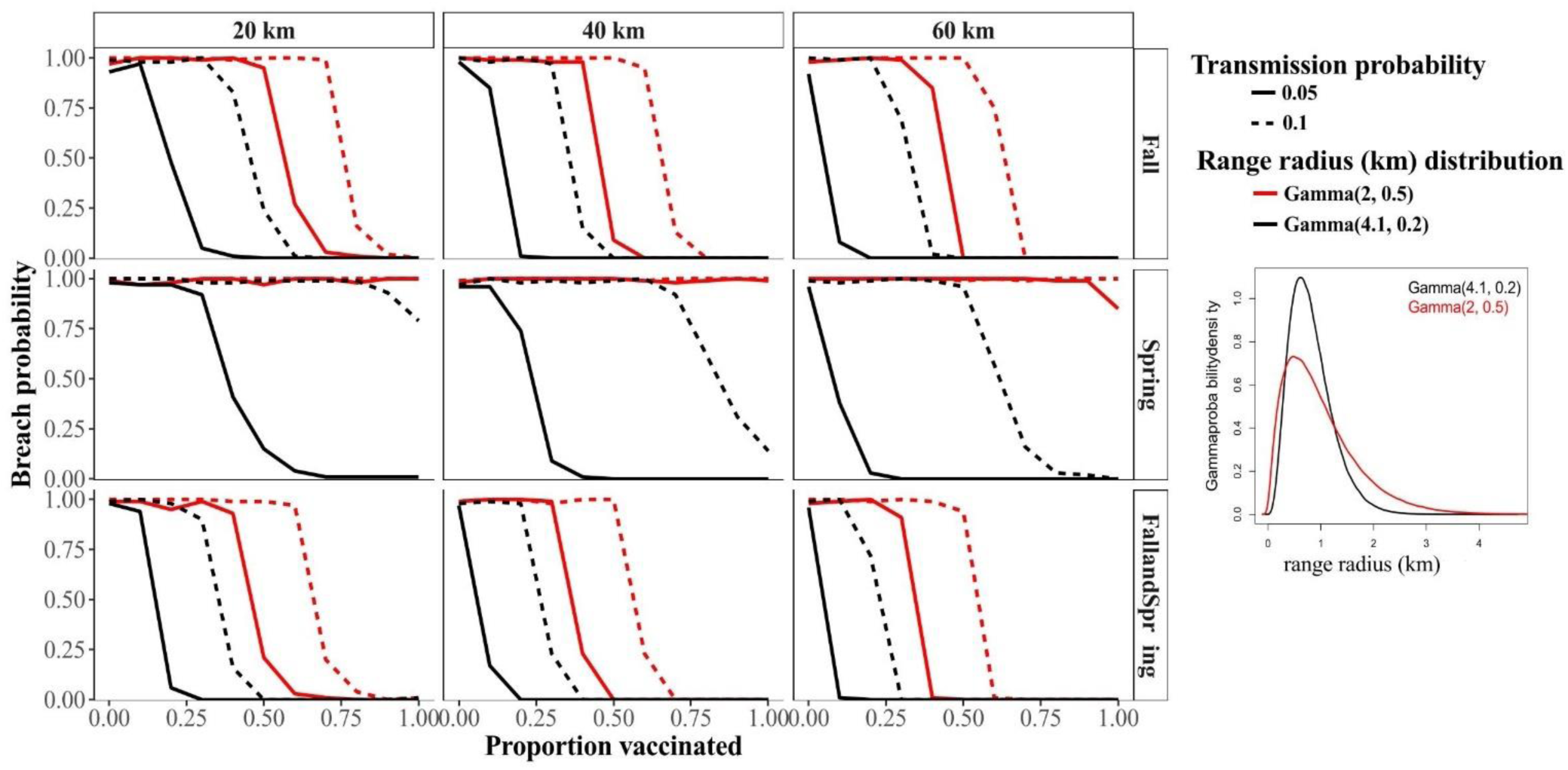
Breach probability given different vaccination strategies. Columns correspond to vaccination zone widths (20km, 40km, or 60km) and rows to vaccination timing (fall, spring, or fall and spring). Line color indicates the weekly home range radius (km) distribution reflecting home range movement. Line type indicates between-group transmission probabilities. Vaccination coverage is indicated along the X-axes. Insert shows gamma probability density functions (with parametrization shape and scale) implemented as weekly-varying home range radius and thus extent.

Variation in the weekly home range radius distribution influenced pathogen persistence probability, which was strongly influenced by both home range size and transmission probability (Fig. 3E-H, Tables S2 & S4). Mean persistence probability was low across much of the parameter space explored in simulations assuming fixed home range areas (mean persistence probability = 0.19, Fig. 3E). For shorter home range radii (<1.5km), variation increased persistence probability by 305-383% relative to fixed radii of the same distance. Within a median radius of 1.5 km, intermediate levels of variation (scale parameter = 1, Fig. 3G) maximized persistence probability relative to more or less variable home range radius distributions, increasing persistence probability by 106% compared to the less variable distributions (Fig. 3F) and by 110% compared to more variable distributions (Fig. 3H). For larger median values of home range radii (1.5-3km), variation increased persistence probability between 110–150% relative to fixed radii of the same length (mean persistence probability = 29% across all simulations implementing variable distributions). In summary, at smaller home range areas, intermediate levels of variation had the highest persistence probability, while at larger home range areas, the effect of variation on persistence weakened considerably relative to simulations with fixed radii.

Annual incidence rates increased with variation in host home range radius distributions relative to most simulations assuming fixed radii (Fig. 3I-L, Fig. 4B, Tables S2 & S5). For shorter home range radii (<1.5km), variation in weekly home range radius distributions increased incidence rates by 167-292%, whereas at longer home range radii (1.5-3km), variation increased incidence rates by 137–192%, when gains in fixed vs. variable home range radii began to diminish. At the highest transmission probabilities, the reverse patterns were observed (Figs. S4B & S5). The relative strength of the effect of variation in home range radii also depended on transmission probability, as seen in Figs 3I-L, and Table S4. Average R_0_ was 0.76 (95% CI (0.697, 0.827)) across 1000 replicate simulations, and 1.72 (95% CI (1.66, 1.77)) for those simulations (444/1000) that did not undergo stochastic fade-out at initiation.

**Figure 4.**
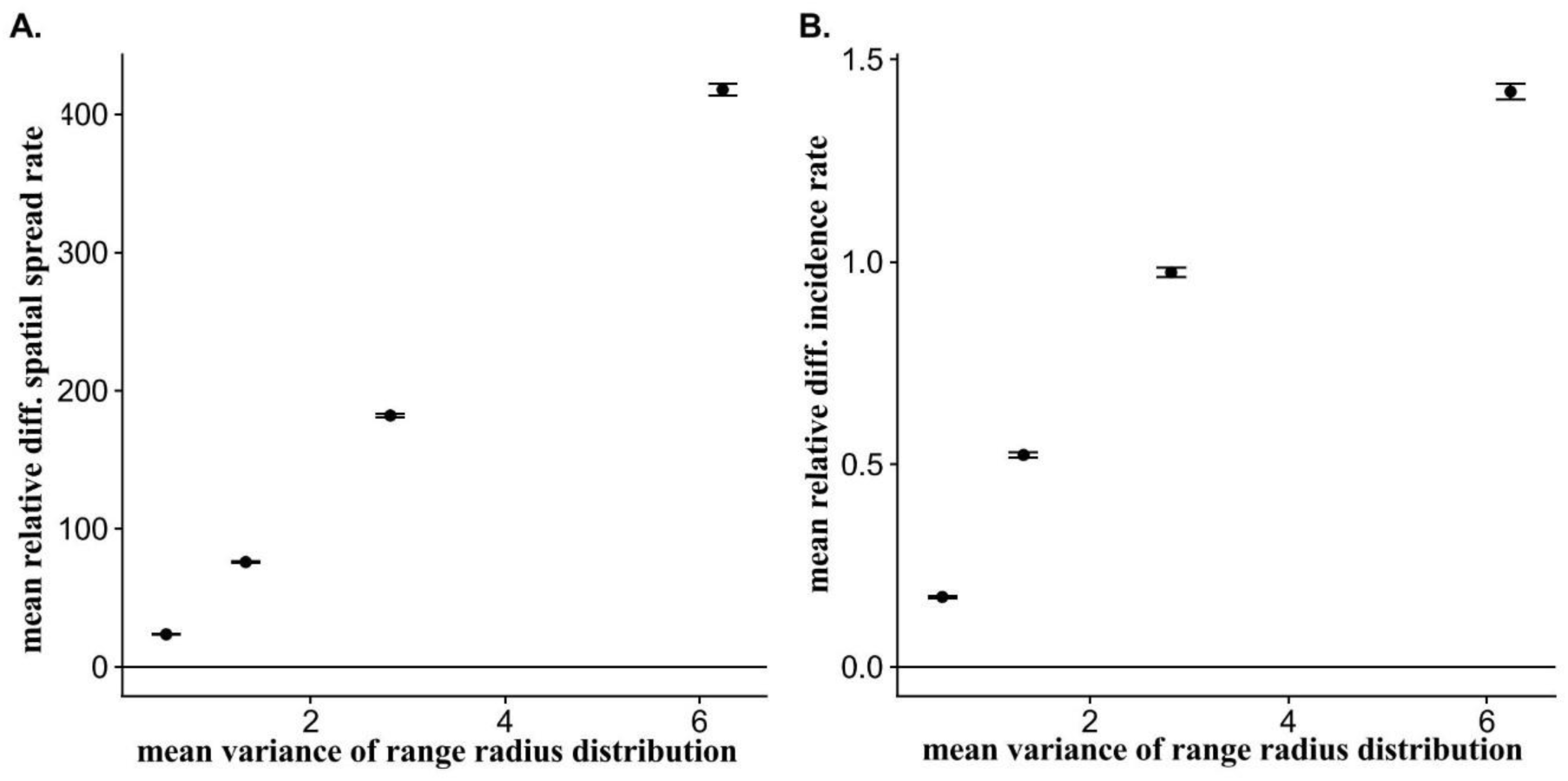
Mean relative difference of fixed vs. variable home range radii in spatial spread rate and incidence rate. **A)** Mean relative difference in spatial spread rates (km/year) between fixed and variable home range radii, calculated as the difference between the variable and fixed home ranges divided by the fixed home range results, plotted against the mean variance of the weekly home range radius (for each of the 4 scale parameters). Points (± 1 SE) are means across all medians and transmission probabilities. **B)** Same as A, but for mean relative difference in incidence rates (new cases/host abundance) between fixed and variable home range radii. Relative differences were evaluated where the median of the gamma distribution = fixed distance moved, in km.

### Effects of home range area variation on vaccination effectiveness

Variation in weekly home range radius distributions strongly influenced ORV effectiveness (Fig. 5). Across all simulations, the home range radius distributions that had higher medians and more variation led to consistent decreases in ORV effectiveness that were driven primarily by increasing variance rather than the median (Fig. S6, Tables S6 & S7). For the fall-only ORV timing, for example, home range radius distributions with more variation decreased relative effectiveness by 54% compared to those with lower variation (red vs. black lines, Fig. 5), while for spring-only ORV timing, higher variation decreased relative effectiveness by 100%. Breach probability increased with transmission probability. In simulations with fall and spring vaccination, for example, doubling the between-group transmission probability decreased ORV effectiveness by 21.7– 41.8% in simulations with higher or lower variation in home range radius distributions, respectively, where all else was held constant.

### Effectiveness of different components of ORV deployment

In simulations with fall ORV timing, vaccination was 40% effective (Table S5), while with spring ORV timing, vaccination was 18.8% effective, when all other conditions were held constant (Fig. 5). When ORV deployment occurred in both fall and spring, it was only marginally more effective than for the fall-only deployment, suggesting diminishing returns with increased vaccination frequency. With the 40km ORV zone, for example, the fall and spring ORV timing increased relative effectiveness by 22% compared to fall-only ORV timing, while for the 60km zone, increasing the frequency of ORV timing yielded a 9.5% relative increase in ORV effectiveness.

ORV effectiveness increased with vaccination coverage, as expected (Fig. 5, Table S7). On average, the minimum coverage required to reduce the probability of breach to zero was 52.8% (range = 0.2 - 1) in simulations where complete reduction was achieved. ORV effectiveness increased with increasing vaccination zone width, but had diminishing returns on breach probability compared to effects of coverage and timing. For example, the 60km ORV area increased relative effectiveness by only 2% compared to the 40km zone, while the 40km zone increased effectiveness by 67% over the 20km zone, when all other conditions were held constant.

## DISCUSSION

We found that small differences in the variation of raccoon home range area had large impacts on the rate of spatial spread of RABV and oral-baited vaccination effectiveness. Our results show that variation in raccoon space use can increase the spread and incidence of RABV, likely by infrequent but substantially longer distance movements of “supermover” individuals (Craft, 2015; White, Forester, & Craft, 2017). We show that interactions between host space use and transmission probability can strongly affect epidemiological processes and vaccination effectiveness, highlighting the need for more information about factors affecting transmission probability and habitat-associated host movement for planning effective control programs.

Variation in host home range area influenced disease dynamics by at least two non-mutually exclusive mechanisms in our model. First, more variable host home ranges increased spatial spread rates because infectious individuals were likely to contact susceptible individuals over longer distances, thus accelerating the spatial spread of RABV to new disease foci. This should have particular importance for species that exhibit heterogeneous population structure and/or social groupings—including lions, jackals, in addition to raccoons—because far-ranging individuals can link spatially or socially isolated groups (Craft, Volz, Packer, & Meyers, 2011; Loveridge & Macdonald, 2001; Russell, Real, & Smith, 2006). Second, variation in host home range area may contribute to host contact heterogeneity. Far-ranging infectious individuals may have more contacts and infect more susceptible individuals, increasing both spread rates and incidence rates, two patterns we report here. Our results suggest that variable host home ranges can drive spatiotemporal variation in contact rates that ultimately affect spatial spread and incidence rates, supporting a growing consensus that variation in host behavior—including host space use—strongly influence wildlife disease dynamics (Dougherty, Seidel, Carlson, Spiegel, & Getz, 2018; Newton et al., 2019; VanderWaal & Ezenwa, 2016).

Our vaccination simulations highlight several key findings for RABV management using ORV zones. First, we found that increases in host home range variation sharply decreased vaccination effectiveness by increasing spatial spread and incidence rates, leading to more frequent vaccination zone breaches at lower to moderate levels of vaccination coverage. Given that a few individuals may disproportionately influence the success or failure of ORV, efforts to better understand drivers of raccoon movement (e.g., conspecific distribution, landscape, disease status) should be a priority, for both infected and uninfected animals. Clinical behaviors of infectious raccoons range from aggressiveness toward conspecifics to paralysis and impaired mobility (Jenkins & Winkler, 1987). Widely roaming infectious individuals (Roscoe et al., 1998) could disproportionately increase disease spread, while paralytic behavior could impede pathogen transmission and slow disease spread. The outcome of pathogen-induced movement behavior on rabies spread may thus depend on the balance of rabid movement behavior among raccoon populations (Reynolds, Hirsch, Gehrt, & Craft, 2015). In uninfected or incubating individuals, individual-level movement behavior can shift in response to disease-induced population declines. For instance, movement patterns and contact rates of red foxes changed as population density decreased following a sarcoptic mange epizootic, leading to increased movement and larger territories (Potts, Harris, & Giuggioli, 2013). Unlike red foxes, however, raccoons exhibit a range of social tolerances including complex seasonally-varying associated and non-associated behaviors, and strict territoriality (Chamberlain & Leopold, 2002; S. D. Gehrt & Fritzell, 1998). Incorporating the interactive effects of pathogen transmission, host-host contact, and host movement behaviors (Hirsch et al., 2013; Prange, Gehrt, & Hauver, 2011; Reynolds et al., 2015) on vaccination effectiveness in this system represents an important avenue for future research.

A second implication for disease management is that bait distribution in fall appears more effective at containing RABV transmission than in spring. Our simulations show seasonal disease dynamics driven by the influx of susceptible juveniles during the synchronous birth pulse in early April to mid-May. Spring vaccination was less effective because it coincides with this birth pulse, when susceptible juveniles are not yet weaned and are unlikely to ingest oral vaccine baits (Fry et al., 2013). Fall vaccination was more effective because susceptible juveniles—who otherwise may have been infectious or incubating the virus—were immunized prior to natal dispersal. Vaccination in both the spring and fall increased effectiveness slightly, but there may be diminishing returns given the relatively small gains in effectiveness and increased implementation costs of a biannual vaccination effort. We note that gains from spring only vaccination or spring and fall together may be greater if protective maternal antibody transmission from vaccinated adult females to young—which we did not model—is prolific in this system. Other components of behavior that we did not model, including breeding and non-breeding contact patterns, may exhibit seasonal variation that could also influence optimal vaccination timing (Reynolds et al., 2015). Our simulations lend support to current ORV timing, but a cost-effectiveness analysis is needed to fully assess the added utility of implementing vaccination twice rather than once per year.

One caveat to this work is that we assumed a homogeneous landscape in our simulations. Landscape heterogeneity can influence the spatial spread of wildlife and plant diseases through scale-dependent effects on host distribution, density, and movement (Meentemeyer, Haas, & Václavík, 2012). At larger spatial scales, topographical features like mountain ranges, rivers, and lakes can influence raccoon movement and partially contain or slow rabies spread among raccoons (Cullingham, Kyle, Pond, Rees, & White, 2009; Smith, Waller, Russell, Childs, & Real, 2005). At smaller spatial scales, spatial heterogeneity resulting from differences in underlying resources influence raccoon foraging behaviors, host movement, and potentially disease processes (Tardy, Massé, Pelletier, & Fortin, 2018; Tardy, Massé, Pelletier, Mainguy, & Fortin, 2014). Importantly, landscape structure can have unexpected consequences on vaccination success when landscape heterogeneity affects host population dynamics and space use. For example, very low vaccination coverage could prevent rabies epizootics that threaten Ethiopian wolves when vaccination is prioritized in dispersal corridors (Haydon et al., 2006). In contrast, low to moderate levels of immunity in raccoons could be counterproductive in landscapes with habitat heterogeneity, because RABV could be perpetuated among weakly connected refuges, igniting future epizootics in neighboring areas (Rees, Pond, Tinline, & Denise, 2013). Considering realistic landscape heterogeneity and mechanistic movement in evaluating disease dynamics and vaccination strategies (e.g. Tracey, Bevins, Vandewoude, & Crooks, 2014; White, Forester, & Craft, 2018) are important directions for future work. A framework that accounts for landscape-driven movement processes would be useful for identifying spatial bait distribution strategies that could increase bait exposure and seroconversion rates, and ultimately, ORV coverage and effectiveness, especially in areas where hand-baiting is used (e.g., urban areas).

A final caveat to this work is that we assumed hosts explored their home range fully and homogenously. This ignores the potential for underlying habitat differences that could affect foraging behaviors, movement, and contact heterogeneity. Recent advances in analytical approaches for studying wildlife space usage, including mechanistic home range movement models that connect underlying movement, resource selection, territoriality, and spatial utilization patterns, are advancing understanding of the behavioral underpinnings of home range animal movement (Börger, Dalziel, & Fryxell, 2008). These methods, in conjunction with parallel advances in approaches using social network theory to investigate host-host contact (Hirsch et al., 2013; Reynolds et al., 2015), offer promise to further elucidate the interacting effects of home range area and host contact structure on disease dynamics and ORV effectiveness, in support of optimizing vaccination strategies for elimination of zoonotic wildlife diseases like RABV.

## ACKNOWLEDGEMENTS

We thank Stacey Elmore for assisting in the literature review for Table S1 and to Carrie Stengel for collecting the raccoon GPS data. Thanks to the USDA APHIS WS National Rabies Management Program, which supported this work. The findings and conclusions in this publication have not been formally disseminated by the USDA and do not represent Agency determination or policy.

## AUTHOR CONTRIBUTIONS

KMP, ATG and EER conceived the ideas and designed methodology, KMP wrote the model code and conducted the simulations, KMP and KMM analyzed the model results, and KMM wrote the first draft of the manuscript. All authors provided critical feedback on the draft and gave final approval for publication.

## SUPPLEMENTAL MATERIALS

**Table S1.** Demographic, movement, and disease parameters used in the vaccination and sensitivity simulations.

| Parameters | Values | References |
| --- | --- | --- |
| **Demographic and movement** |  |  |
| Longevity (number of years before natural death occurs) | $\sim \text{EXP}(\lambda)$, $\lambda = 0.3$; mean = 3 years | (S. D. Gehrt & Prange, 2007; Johnson, 1970; Nowak, 1999; Prange, Gehrt, & Wiggers, 2003) |
| Age at reproductive maturity (minimum age females will conceive) | 13 months | (Lotze & Anderson, 1979) |
| Annual conception probability (by age class of females) | 0.93 ($\geq 1.5$ years),<br>0.54 ($< 1.5$ years) | (Prange et al., 2003) |
| Litter size (number of offspring per litter) | $\sim \text{NORM}(\mu, \sigma^2)$; $\mu = 4$ kits, $\sigma^2 = 1$ | (Fritzell, Hubert Jr, Meyen, & Sanderson, 1985; Ritke, 1990; G. C. Sanderson & Hubert, 1981) |
| Gestation period | 9 weeks | (G. Sanderson & Nalbandov, 1973) |
| Maximum group size | 10 individuals | expert opinion |
| Grid-cell carrying capacity | 15 raccoons/km<sup>2</sup> | (Kennedy, Nelson, Weckerly, & Sugg, 1991; Moore & Kennedy, 1985; Sonenshine & Winslow, 1972; Urban, 1970) |
| Age that male dyads dissolve and males become independent | 1.5 years | (S. Gehrt & Fritzell, 1998a); expert opinion |
| Dispersal age of males from the group | $\sim \text{POI}(\lambda)$, $\lambda = 10$ months | (S. Gehrt & Fritzell, 1998b) |
| Dispersal distance (males for natal dispersal, both males and females for oversized groups) | $\sim \text{WEIBULL}(\lambda, k)$, $\lambda = 1.5$, $k = 0.3$<br>median = 0.44 km, mean = 13.6 km<br>variance = 5527 | (Cullingham et al., 2008; Dharmarajan, Beasley, Fike, & Rhodes, 2009) |
| Weekly distance moved in km (relative to home range centroid) | $\sim \text{GAM}(k, \theta)$, 1) $k = 4.1$, $\theta = 0.2$, 2) $k = 2$, $\theta = 0.25$ | J. Doe, unpublished data |
| **Disease** |  |  |
| Incubation period | $\sim \text{POI}(\lambda)$, $\lambda = 4$ weeks | (Rees, Pond, Phillips, & Murray, 2008; Tinline, Rosatte, & MacInnes, 2002) |
| Infectious period | 1 week | (Hanlon, Niezgoda, & Rupprecht, 2007) |
| Recovery rate/probability | 0.1 | expert opinion |
| Probability of disease-induced mortality | 1 | (Hanlon et al., 2007; Rupprecht et al., 1986) |
| Within-group transmission probability | 0.5 | NA |
| Between-group transmission probability | 0.001 - 0.5 | NA |

**Figure S1.**
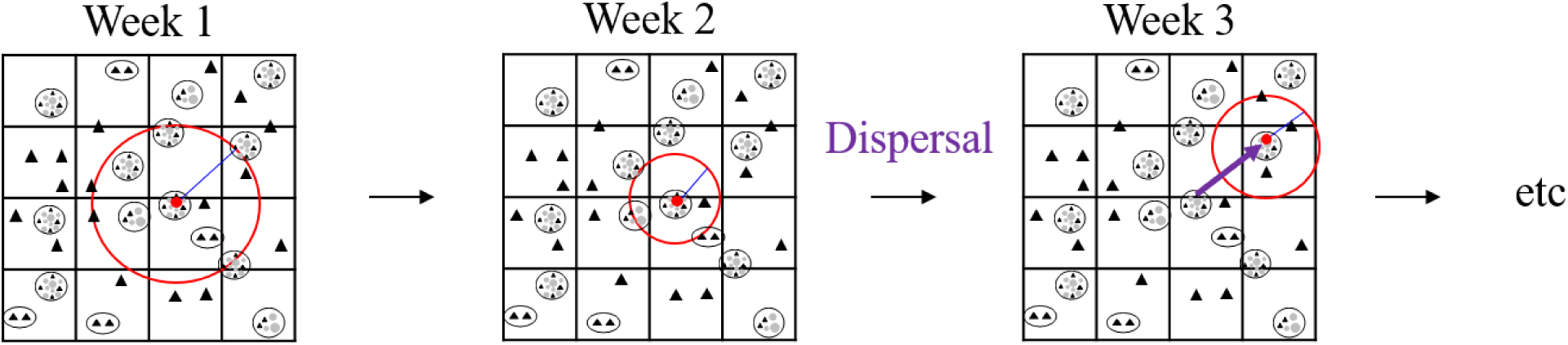
Host movement and contact structure. Schematic representing weekly home range movement and resulting host-host contact within the gridded simulated landscape. Filled triangles are males, filled circles are females, and family groups and male dyads are enclosed by black circles. Individuals move a weekly-varying distance (the radius of the circular home range; shown by a blue line) that originates from the family group home centroid and is drawn from a gamma distribution. The red circle represents the home range explored by one individual host on a particular week. Natal dispersal (shown in purple) represents longer distance movements undertaken primarily by males, with dispersal distance drawn from a Weibull distribution. Contact among infectious and susceptible individuals and pathogen transmission (if applicable; see main text for details) occurs within the home range (red circle) of each infectious individual at every weekly time step.

**Figure S2.**
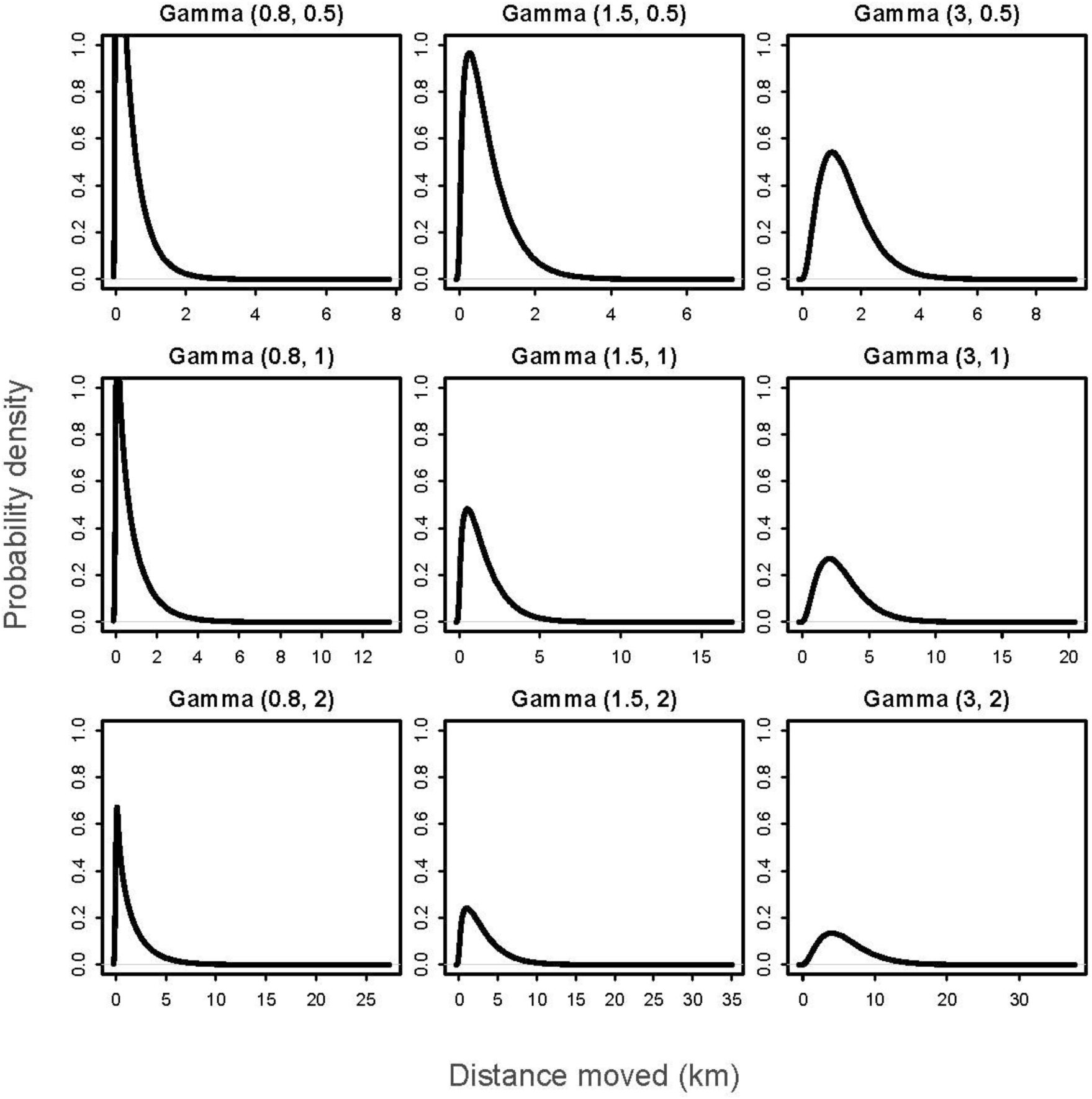
Gamma probability density functions. Example probability density functions for gamma distributions over the range of parameters used in the simulations with variable weekly distance moved. At each time step, each individual moved a distance drawn from a particular gamma distribution parameterized with a shape and scale parameter (Gamma (shape, scale)).

**Figure S3.**
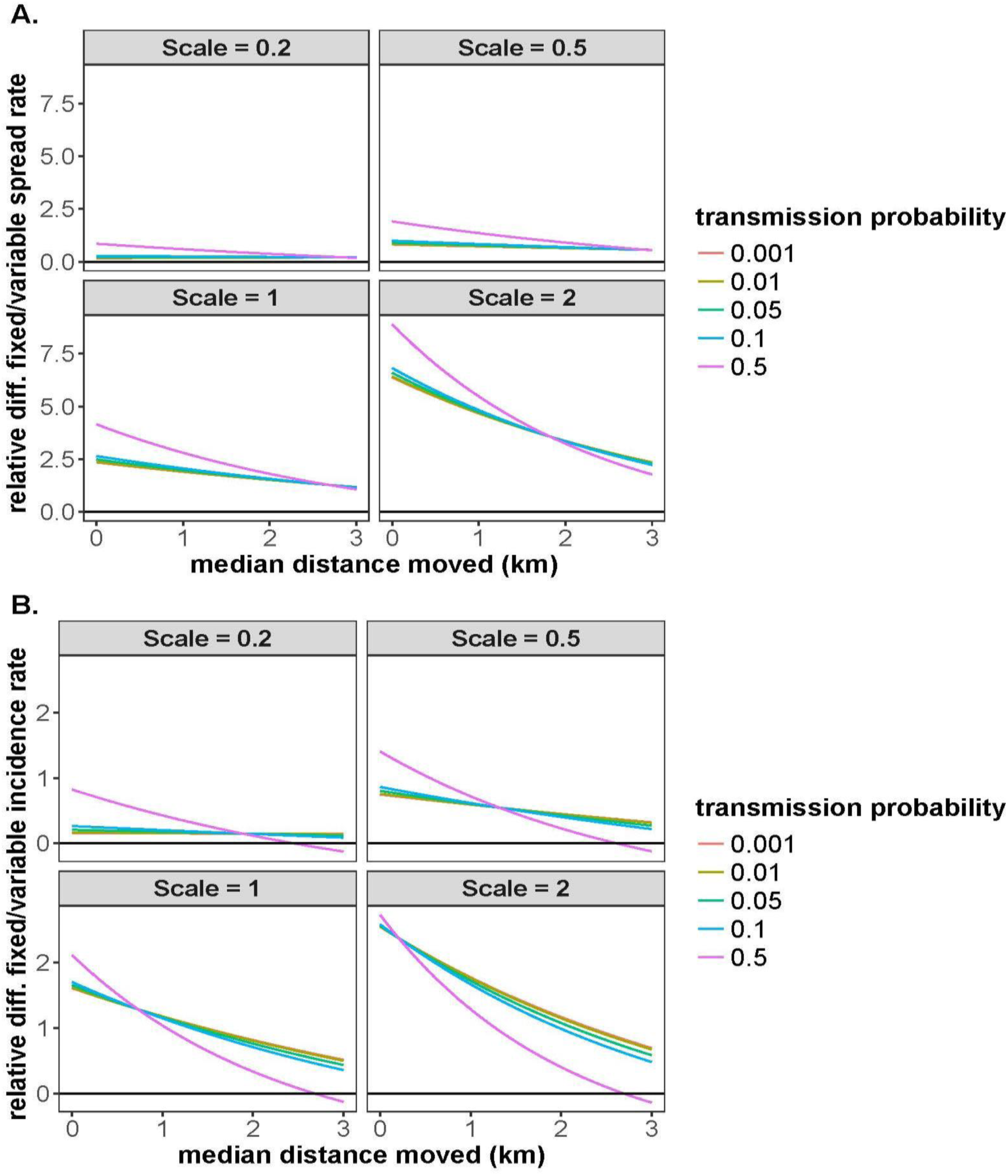
Relative difference of fixed vs. variable movement in spatial spread rate and incidence rate. A) Relative difference in predicted spatial spread rates (km/year) between fixed and variable movement simulations plotted against the median (distance moved) of the gamma-distributed weekly movement random variable. B) Relative difference in predicted incidence rates (annual new cases/annual mean population size) between fixed and variable movement simulations plotted against the median (distance moved) of the gamma-distributed weekly movement random variable. Relative differences were calculated as the difference between the predicted variable movement simulation response and predicted fixed movement simulation response divided by the predicted fixed movement simulation response, evaluated where the median of the gamma movement distribution = fixed distance moved in km. Scale refers to the scale parameter of the weekly gamma movement distribution implemented in the simulations which controls the spread of the distribution. Variance in the weekly distance moved by individuals increases with increasing values of the scale parameter. The black line along 0 indicates no difference between fixed and variable conditions. Values above 0 indicate the relative increase in response variables due to variation in the movement distribution.

**Figure S4.**
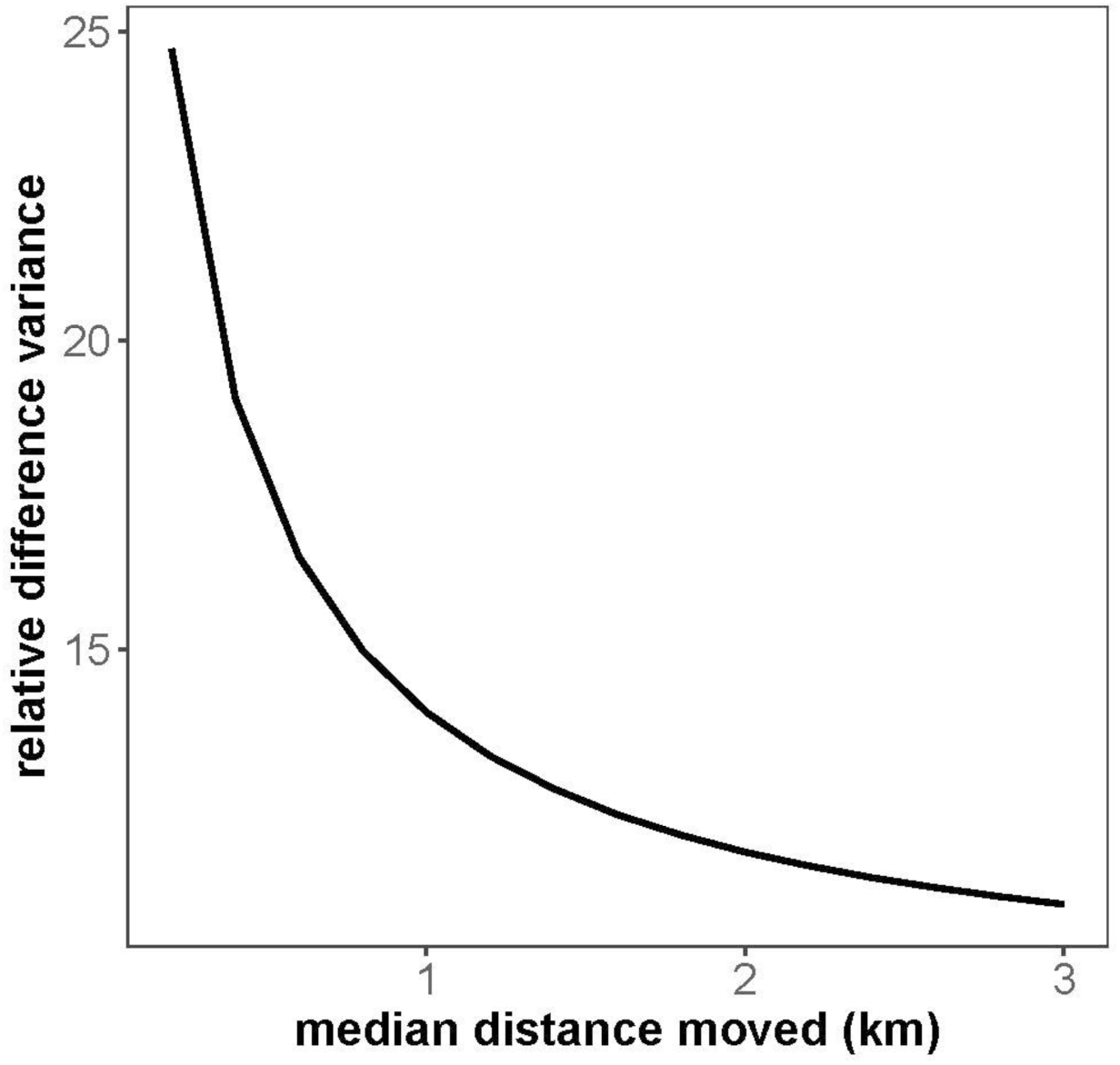
Relative difference in variance of high and low variance movement distributions used in simulations. Relative difference in the weekly host distribution variance between simulations with the highest variance (where the scale parameter of the weekly host movement gamma distribution = 2) vs. the lowest variance (where the scale parameter of the gamma distribution = 0.2) plotted against the median distance moved (km) of the weekly movement gamma distribution. Relative difference was calculated as the difference in the variance of the gamma distribution in high and low variance movement distributions divided by the variance in the low variance movement distribution. This shows that at lower median distances moved, the variance associated with the more variable movement distributions is higher relative to less variable movement distributions, but this difference diminishes as median distance moved increases.

**Figure S5.**
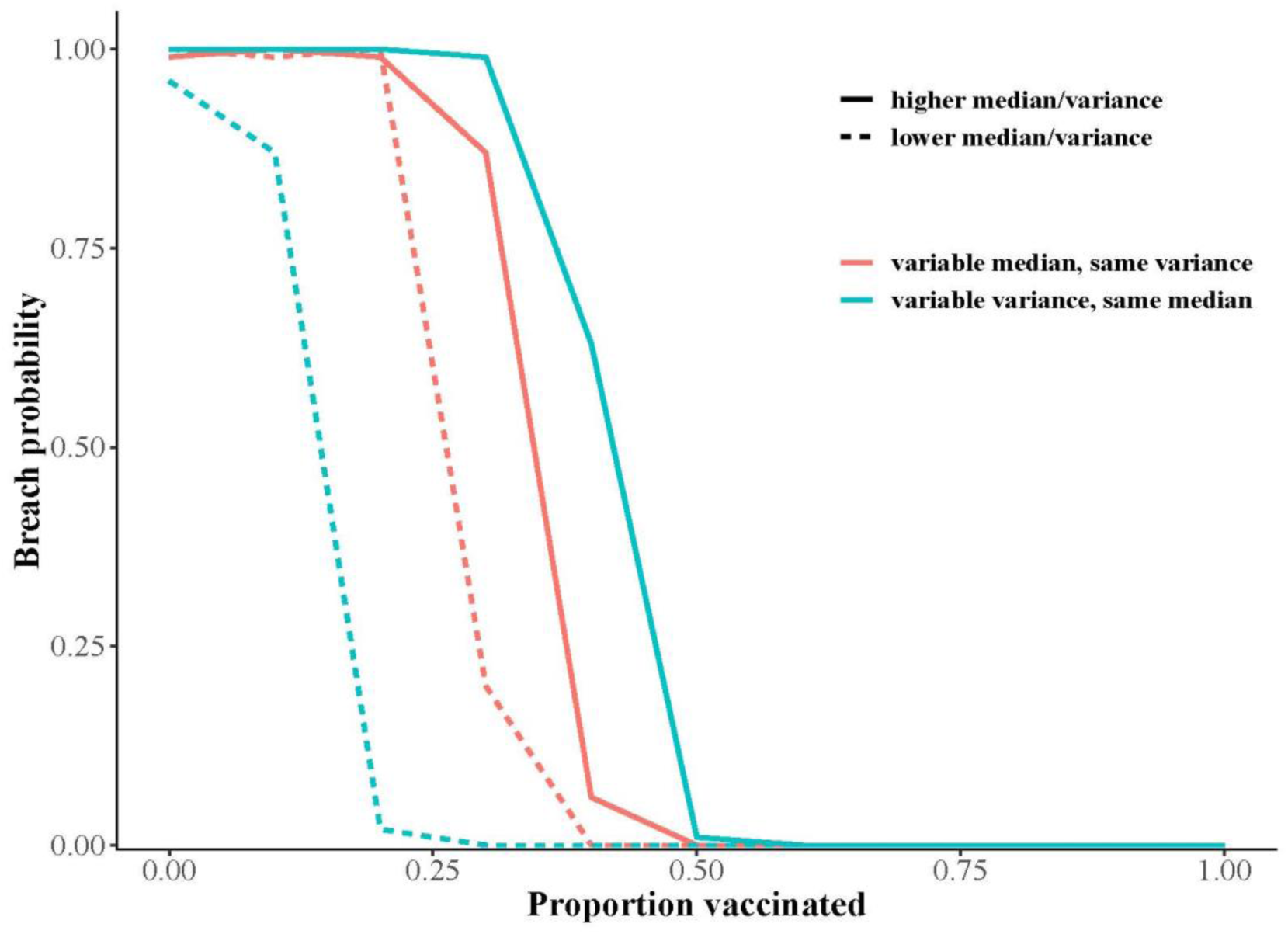
Breach probability given different weekly movement distributions. Rabies breach probability plotted against vaccination coverage for vaccination simulations. Line color represents weekly movement distributions with variable medians (0.75km, 0.87km) but similar variance (0.3), or variable variance (0.49, 0.17) but similar medians (0.8km). Line type indicates higher and lower values for median or variance. Breach probability is defined as the proportion of 100 replicate simulations in which the vaccination zone was breached.

**Figure S6.**
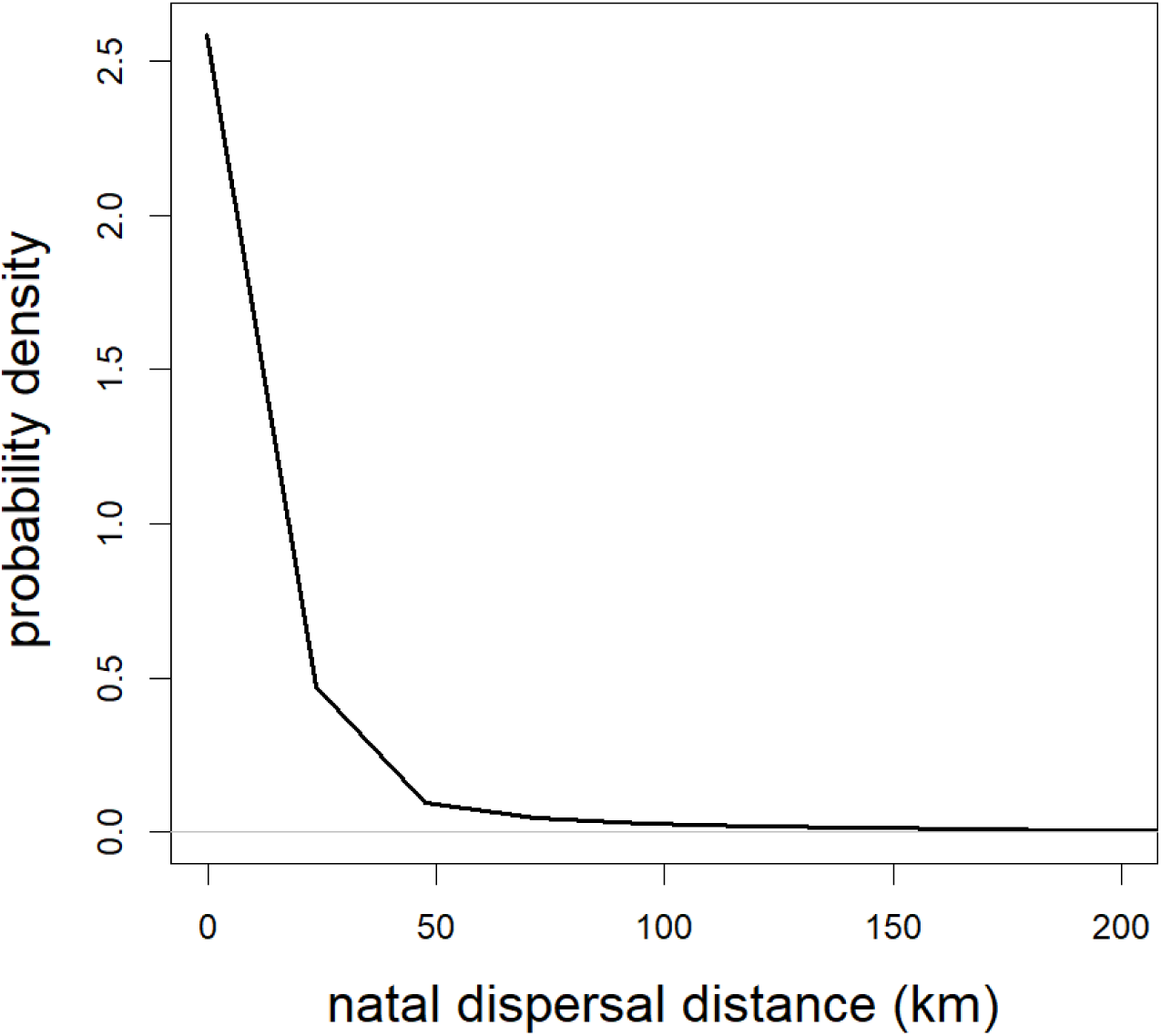
Weibull probability density function. Dispersal distances related to natal dispersal and to avoid over-crowding in the model were modeled as a Weibull random variable. Dispersal distances were drawn from a Weibull distribution (shown here) that was parameterized with a scale and shape parameter (Weibull(λ, k), λ = 1.5, k = 0.3), with a median and mean dispersal distance of 0.44 km and 13.6 km, respectively, and a variance of 5527.

**Figure S7.**
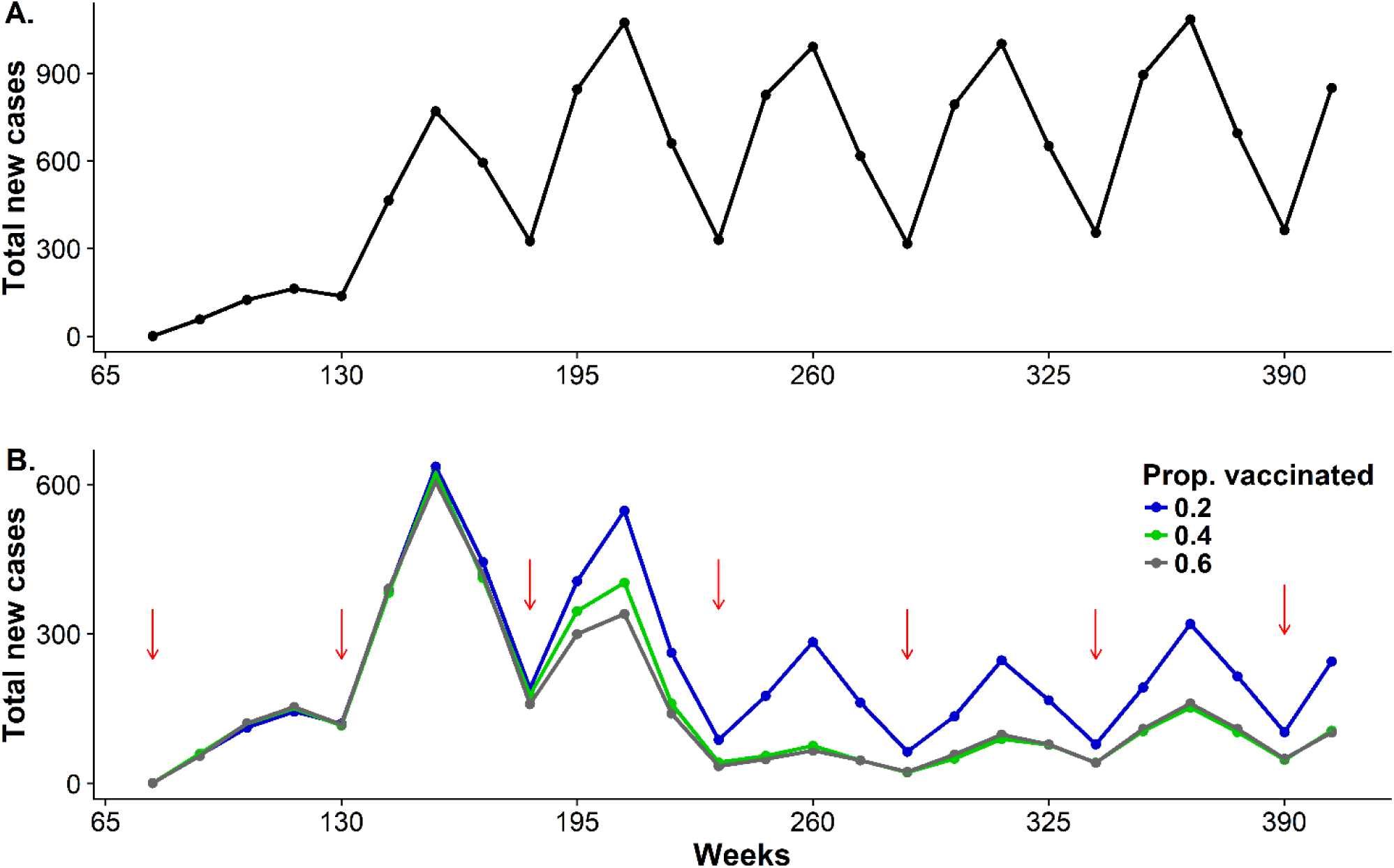
Total cases with and without vaccination. **A)** Disease dynamics in the absence of vaccination. Points are the mean total new cases at a quarterly time step (13 weeks per time step), estimated as the average of 100 simulations with the same parameter values. **B)** Points are mean total new cases for three vaccination coverage scenarios (0.2, 0.4, and 0.6 of population vaccinated). Red arrows are the quarterly time step in which vaccines were deployed (fall vaccination shown here). In both A. and B., the simulated vaccination zone width was 40km, between-group transmission probability was 0.05, and weekly movement distribution was a gamma distribution where shape = 4.1 and scale = 0.2. Standard deviation was excluded to more easily visualize patterns.

## I. STATISTICAL ANALYSES RESULTS

### A. Statistical analysis of sensitivity simulations

**Response variables:**

> **Annual spatial spread rate (km/year);** continuous (annual linear distance traveled per year during which disease was present at an incidence rate ≥ 0.001)
>
> **Persistence probability;** binary (whether or not there were rabies cases in the last time step of simulation)
>
> **Annual incidence rate;** continuous (average annual number of new rabies cases/annual maximum population size across years in which disease was present)

Main effects:
| Variable/Factor | Levels |
| --- | --- |
| **Fixed distance moved/median of movement distribution** | 0.2 – 3 km in 0.2 increments |
| **Shape parameter of gamma movement distribution** | 0.2, 0.5, 1, 2 |
| **Between-group transmission probability** | 0.001 to 0.5 in logarithmic intervals |

**Table S2.** Results of AIC analysis for generalized linear models of: 1) annual spatial spread rate and 2) annual incidence rate, both with a gamma distribution and a log link function, and 3) persistence probability with a binomial distribution and a logit link function. Results shown for simulations with both constant and stochastic home range movement. Main effects were included in all models where 2-way interactions are indicated. K is the number of parameters. Aikaike weights were calculated to provide a measure of model selection uncertainty among fitted models (Burnham & Anderson, 2002). Asterisk indicates model with an Aikaike weight of 1.

| Response variable | Movement model | Model specification | K | log likelihood | AICc | Delta AIC |
| --- | --- | --- | --- | --- | --- | --- |
| Spatial spread rate | Fixed | *fixed distance moved x trans. prob. | 4 | -75743.38 | 151496.8 | 0 |
|  |  | fixed distance moved + trans. prob. | 3 | -75784.53 | 151577.1 | 80.30725 |
|  |  | Transmission probability | 2 | -82282.08 | 164570.2 | 13073.39249 |
|  |  | fixed distance moved | 2 | -79320.55 | 158647.1 | 7150.34216 |
|  |  | Intercept only | 1 | -83737.75 | 167479.5 | 15982.74677 |
|  | Variable | *All main effects and 2-way interactions | 13 | -392476.7 | 784981.5 | 0 |
|  |  | Median x scale + trans. prob | 9 | -392533.1 | 785086.2 | 104.726 |
|  |  | Scale x trans. prob. + median | 9 | -394437.8 | 788895.6 | 3914.16 |
|  |  | Median x trans. prob + scale | 7 | -394441.8 | 788899.6 | 3918.158 |
|  |  | Trans. prob x scale | 8 | -415863.4 | 831744.8 | 46763.375 |
|  |  | Median x trans. prob | 4 | -409855 | 819720.1 | 34738.633 |
|  |  | Median x scale | 8 | -408415.6 | 816849.3 | 31867.846 |
|  |  | Median + scale + trans. prob | 6 | -394445.7 | 788905.4 | 3923.94 |
|  |  | Scale + trans. prob | 5 | -415867.3 | 831746.5 | 46765.097 |
|  |  | Median + trans. prob | 3 | -409856.6 | 819721.2 | 34739.785 |
|  |  | Median + scale | 5 | -409603.1 | 819218.2 | 34236.76 |
|  |  | Scale | 4 | -423523 | 847056 | 62074.541 |
|  |  | Transmission probability | 2 | -422776.9 | 845559.8 | 60578.37 |
|  |  | Median | 2 | -419990.6 | 839987.3 | 55005.841 |
|  |  | Intercept only | 1 | -429034.1 | 858072.3 | 73090.832 |
| Incidence rate | Fixed | *fixed distance moved x trans. prob. | 4 | -11247.86 | 22503.72 | 0 |
|  |  | fixed distance moved + trans. prob. | 3 | -11903.1 | 23812.2 | 1308.479 |
|  |  | Transmission probability | 2 | -12755.8 | 25515.6 | 3011.876 |
|  |  | fixed distance moved | 2 | -12586.91 | 25177.83 | 2674.105 |
|  |  | Intercept only | 1 | -13404.09 | 26810.18 | 4306.461 |
|  | **Variable** | *All main effects and 2-way interactions | 13 | -50154.35 | 100334.7 | 0 |
|  |  | Median x scale + trans. prob | 9 | -58006.74 | 116031.5 | 15696.787 |
|  |  | Scale x trans. prob. + median | 7 | -51887.8 | 103789.6 | 3454.898 |
|  |  | Median x trans. prob + scale | 9 | -57490.31 | 114998.6 | 14663.932 |
|  |  | Trans. prob x scale | 8 | -58071.49 | 116159 | 15824.283 |
|  |  | Median x trans. prob | 4 | -51993.64 | 103995.3 | 3660.592 |
|  |  | Median x scale | 8 | -59269.4 | 118554.8 | 18220.103 |
|  |  | Median + scale + trans. prob | 6 | -58307.5 | 116627 | 16292.308 |
|  |  | Scale + trans. prob | 5 | -58372.04 | 116754.1 | 16419.378 |
|  |  | Median + trans. prob | 3 | -58401.53 | 116809.1 | 16474.359 |
|  |  | Median + scale | 5 | -60062.01 | 120134 | 19799.322 |
|  |  | Scale | 4 | -60124.56 | 120257.1 | 19922.431 |
|  |  | Transmission probability | 2 | -58465.93 | 116935.9 | 16601.165 |
|  |  | Median | 2 | -60153.16 | 120310.3 | 19975.621 |
|  |  | Intercept only | 1 | -60215.59 | 120433.2 | 20098.494 |
| **Persistence prob.** | **Fixed** | *fixed distance moved x trans. prob. | 4 | 4297.39 | -8584.777 | 0 |
|  |  | fixed distance moved + trans. prob. | 3 | 4236.4 | -8464.797 | 119.9794 |
|  |  | trans. prob | 2 | 2498.8 | -4991.599 | 3593.1779 |
|  |  | fixed distance moved | 2 | 3359.685 | -6713.369 | 1871.4074 |
|  |  | Intercept only | 1 | 2123.488 | -4242.976 | 4341.8009 |
|  | **Variable** | *All main effects and 2-way interactions | 13 | -1527.572 | 3083.148 | 0 |
|  |  | Median x scale + trans. prob | 9 | -1895.851 | 3811.706 | 104.726 |
|  |  | Median x trans. prob + scale | 9 | -2061.625 | 4139.252 | 3914.16 |
|  |  | Scale x trans. prob. + median | 7 | -2287.333 | 4594.669 | 3918.158 |
|  |  | Trans. prob + scale | 5 | -7325.597 | 14669.196 | 46763.375 |
|  |  | Median x trans. prob | 4 | -2837.068 | 5684.137 | 34738.633 |
|  |  | Median x scale | 8 | -4339.501 | 8697.004 | 31867.846 |
|  |  | Median + scale + trans. prob | 6 | -2306.647 | 4627.295 | 3923.94 |
|  |  | Scale + trans. prob | 5 | -7333.577 | 14679.155 | 46765.097 |
|  |  | Median + trans. prob | 3 | -3048.917 | 6105.834 | 34739.785 |
|  |  | Median + scale | 5 | -4613.832 | 9239.666 | 34236.76 |
|  |  | Scale | 4 | -8523.421 | 17056.843 | 62074.541 |
|  |  | Trans. prob | 2 | -7599.468 | 15204.937 | 60578.37 |
|  |  | Median | 2 | -5138.863 | 10283.727 | 55005.841 |
|  |  | Intercept only | 1 | -8737.917 | 17479.835 | 73090.832 |

**Table S3.** Parameter estimation of best-supported annual spatial spread rate model using Akaike Information Criterion (AIC; 2 points). Analyses support Figure 3A-D.

| Fixed distance moved | Estimate | SE | t value | P value |
| --- | --- | --- | --- | --- |
| Intercept | -13.376 | 0.506 | -26.432 | <0.0001 |
| Fixed distance moved | 15.077 | 0.156 | 96.710 | <0.0001 |
| Transmission probability | 3.170 | 2.347 | 1.351 | 0.18 |
| Trans. prob. x distance | 43.327 | 0.772 | 56.087 | <0.0001 |

| Stochastic distance moved | Estimate | SE | t value | P value |
| --- | --- | --- | --- | --- |
| Intercept | 1.845 | 0.011 | 167.154 | <0.0001 |
| Median of gamma distribution | 0.545 | 0.003 | 160.637 | <0.0001 |
| Scale 0.5 | 0.442 | 0.014 | 30.634 | <0.0001 |
| Scale 1 | 1.051 | 0.014 | 75.159 | <0.0001 |
| Scale 2 | 1.842 | 0.014 | 135.424 | <0.0001 |
| Transmission probability | 2.837 | 0.036 | 79.584 | <0.0001 |
| Median x scale 0.5 | -0.062 | 0.004 | -14.060 | <0.0001 |
| Median x scale 1 | -0.155 | 0.004 | -35.675 | <0.0001 |
| Median x scale 2 | -0.275 | 0.004 | -64.599 | <0.0001 |
| Median x trans. prob. | -0.066 | 0.009 | -7.039 | <0.0001 |
| Scale 0.5 x trans. prob. | 0.019 | 0.036 | 0.523 | 0.601 |
| Scale 1 x trans. prob. | -0.060 | 0.036 | -1.660 | 0.097 |
| Scale 2 x trans. prob. | -0.336 | 0.036 | -9.337 | <0.0001 |

n = number observations = 46,402
K= number of parameters = 13
R<sup>2</sup> = 0.59

**Table S4.** Parameter estimation of best-supported pathogen persistence probability model using Akaike Information Criterion (AIC; 2 points). Analyses support Figure 3E-H.

| **Fixed distance moved** | **Estimate** | **SE** | **t value** | **P value** |
| --- | --- | --- | --- | --- |
| Intercept | -2.573 | 0.043 | -59.196 | <0.0001 |
| Fixed distance moved | 0.724 | 0.014 | 50.085 | <0.0001 |
| Transmission probability | 5.600 | 0.302 | 18.529 | <0.0001 |
| Trans. prob. x distance | -7.449 | 0.220 | -33.911 | <0.0001 |
| n = number observations = 15,000<br>K = number of parameters = 4<br>R<sup>2</sup> = 0.184 |  |  |  |  |
| **Stochastic distance moved** | **Estimate** | **SE** | **t value** | **P value** |
| Intercept | -2.087 | 0.038 | -55.345 | <0.0001 |
| Median of gamma distribution | 0.697 | 0.013 | 54.587 | <0.0001 |
| Transmission probability | 14.342 | 0.284 | 50.572 | <0.0001 |
| Scale 0.5 | 0.719 | 0.050 | 14.386 | <0.0001 |
| Scale 1 | 1.523 | 0.049 | 31.148 | <0.0001 |
| Scale 2 | 2.176 | 0.049 | 44.022 | <0.0001 |
| Median x trans. prob. | -13.993 | 0.168 | -83.305 | <0.0001 |
| Scale 0.5 x trans. prob. | 2.581 | 0.321 | 8.032 | <0.0001 |
| Scale 1 x trans. prob. | -0.332 | 0.322 | -1.031 | 0.303 |
| Scale 2 x trans. prob. | -10.371 | 0.352 | -29.447 | <0.0001 |
| Median x scale 0.5 | -0.235 | 0.017 | -14.100 | <0.0001 |
| Median x scale 1 | -0.505 | 0.016 | -30.597 | <0.0001 |
| Median x scale 2 | -0.752 | 0.017 | -44.557 | <0.0001 |
| n = number observations = 60,000<br>K = number of parameters = 13<br>R<sup>2</sup> = 0.186 |  |  |  |  |

**Table S5.** Parameter estimation of best-supported annual incidence rate model using Akaike Information Criterion (AIC; 2 points). Analyses support Figure 3I-L.

| **Fixed distance moved** | **Estimate** | **SE** | **t value** | **P value** |
| --- | --- | --- | --- | --- |
| Intercept | -3.044 | 0.018 | 169.934 | <0.0001 |
| Fixed distance moved | 0.554 | 0.006 | 98.856 | <0.0001 |
| Transmission probability | 3.947 | 0.086 | 45.991 | <0.0001 |
| Trans. prob. x distance | -0.582 | 0.029 | -20.391 | <0.0001 |
| n = number observations = 9,732 |  |  |  |  |
| K = number of parameters = 4 |  |  |  |  |
| R<sup>2</sup> = 0.34 |  |  |  |  |
| **Stochastic distance moved** | **Estimate** | **SE** | **t value** | **P value** |
| Intercept | -2.716 | 0.012 | -227.10 | <0.0001 |
| Median of gamma distribution | 0.489 | 0.004 | 131.264 | <0.0001 |
| Scale 0.5 | 0.414 | 0.016 | 26.099 | <0.0001 |
| Scale 1 | 0.815 | 0.015 | 52.872 | <0.0001 |
| Scale 2 | 1.122 | 0.015 | 74.761 | <0.0001 |
| Transmission probability | 3.352 | 0.040 | 83.465 | <0.0001 |
| Median x scale 0.5 | -0.091 | 0.005 | -18.434 | <0.0001 |
| Median x scale 1 | -0.179 | 0.005 | -37.039 | <0.0001 |
| Median x scale 2 | -0.244 | 0.005 | -51.462 | <0.0001 |
| Median x trans. prob. | -0.485 | 0.011 | -46.000 | <0.0001 |
| Scale 0.5 x trans. prob. | -0.274 | 0.041 | -6.670 | <0.0001 |
| Scale 1 x trans. prob. | -0.559 | 0.041 | -13.626 | <0.0001 |
| Scale 2 x trans. prob. | -0.813 | 0.041 | -19.912 | <0.0001 |
| n = number observations = 47,378 |  |  |  |  |
| K = number of parameters = 13 |  |  |  |  |
| R<sup>2</sup> = 0.3 |  |  |  |  |

### B. Statistical analysis of vaccination simulations

**Response:**

Binary (whether or not rabies breached the vaccination zone)

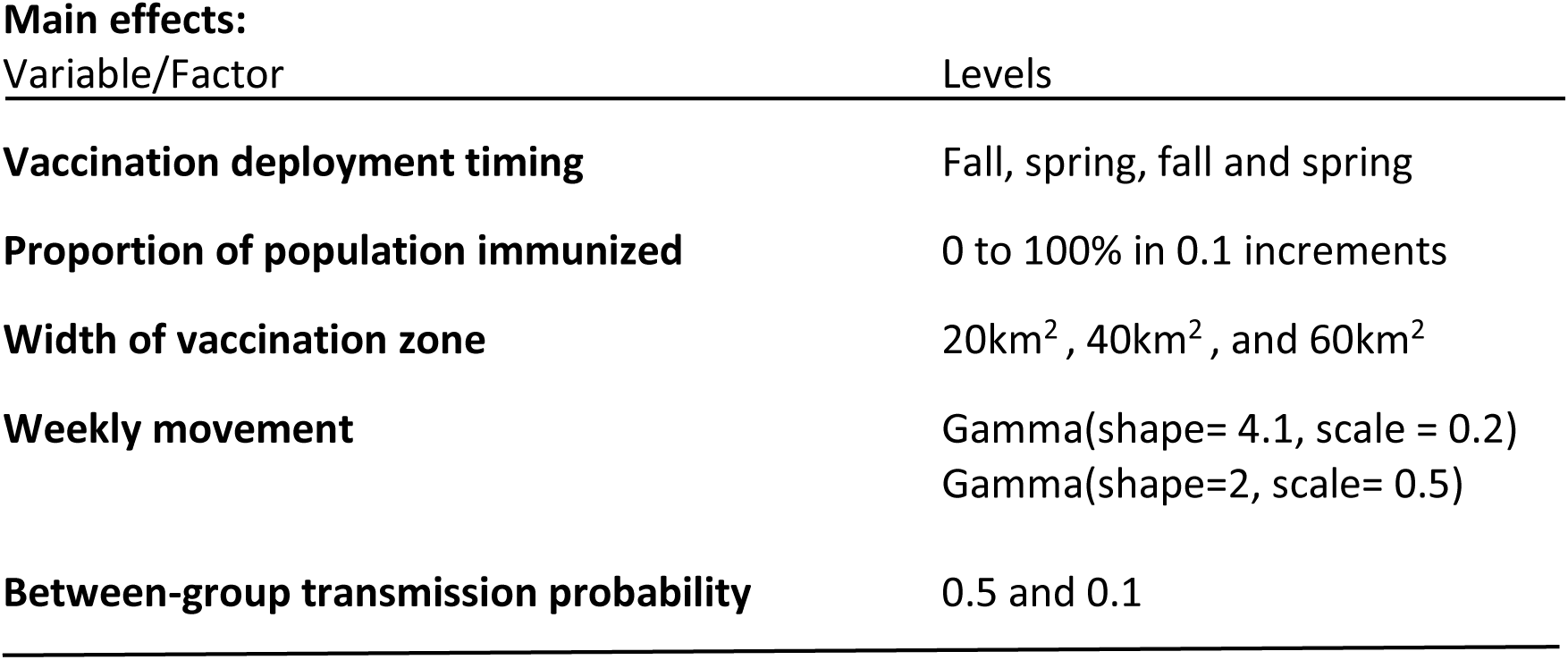

**Table S6.** Results of AIC analysis for generalized linear models of vaccination zone breach probability with a binomial distribution and a logit link function for vaccination simulations. Main effects were included in all models where 2-way interactions are indicated. K is the number of parameters in the model. K is the number of parameters in a model. The top ten best-supported models are shown. Aikaike weights were calculated to provide a measure of model selection uncertainty among fitted models (Burnham & Anderson, 2002). Asterisk indicates model with an Aikaike weight of 1.

| Model specification | K | log likelihood | AICc | Delta AIC | AIC weight |
| --- | --- | --- | --- | --- | --- |
| All main effects and 2-way interactions | 27 | -3700.075 | 7454.188 | 0 | 1 |
| All interactions except prop. vaccinated: trans. prob. | 26 | -3722.473 | 7496.982 | 42.79354 | 0 |
| All interactions except timing x trans. prob. | 25 | -3728.677 | 7507.387 | 53.19927 | 0 |
| All interactions except prop. vaccinated x trans. prob. & timing x trans. prob. | 24 | -3730.56 | 7509.151 | 54.96322 | 0 |
| All interactions except area x trans. prob. | 25 | -3732.351 | 7514.735 | 60.54698 | 0 |
| All interactions except area x trans. prob. & timing x trans. prob. | 24 | -3738.66 | 7525.351 | 71.16281 | 0 |
| All interactions except prop. vaccinated x trans. prob. & timing x trans. prob. & areas x trans. prob. | 22 | -3747 | 7538.026 | 83.83756 | 0 |
| All interactions except area x trans. prob. & timing x trans. prob. | 23 | -3746.999 | 7540.026 | 85.83786 | 0 |
| All interactions except area x timing | 23 | -3753.471 | 7552.971 | 98.78287 | 0 |
| All interactions except area x trans. prob. & area x timing | 21 | -3759.986 | 7561.995 | 107.8067 | 0 |

**Table S7.** Parameter estimation of the best-supported model of breach probability using Akaike Information Criterion (AIC; 2 points), which includes all main effects and all possible two-way interactions. Simulation data was fit to generalized linear models with a binomial distribution and a logit link. Analyses support Figure 5.

| Model term | Estimate | SE | t value | P value |
| --- | --- | --- | --- | --- |
| Intercept | 3.23266 | 0.18231 | 17.732 | <0.0001 |
| shape_2 | 10.17199 | 0.37365 | 27.223 | <0.0001 |
| transmission probability_0.1 | 9.40278 | 0.3173 | 29.634 | <0.0001 |
| proportion vaccinated | -45.7251 | 1.14096 | -40.076 | <0.0001 |
| timing_spring | -1.3584 | 0.21207 | -6.405 | <0.0001 |
| timing_spring+fall | -2.32445 | 0.20602 | -11.283 | <0.0001 |
| barrier area_20km | 4.70812 | 0.23768 | 19.808 | <0.0001 |
| barrier area_40km | 2.18564 | 0.19935 | 10.964 | <0.0001 |
| shape_2 x trans. prob._0.1 | -5.14293 | 0.27737 | -18.542 | <0.0001 |
| shape_2 x prop. vaccinated | 13.78053 | 0.64275 | 21.44 | <0.0001 |
| shape_2 x timing_spring | -5.15968 | 0.42305 | -12.196 | <0.0001 |
| shape_2 x timing_spring+fall | 1.85079 | 0.44156 | 4.191 | <0.0001 |
| shape_2 x area_20km | -4.31862 | 0.28276 | -15.273 | <0.0001 |
| shape_2 x area_40km | -2.30984 | 0.28159 | -8.203 | <0.0001 |
| trans. prob._0.1 x prop vaccinated | 3.98704 | 0.56487 | 7.058 | <0.0001 |
| trans. prob._0.1 x timing_spring | -2.38452 | 0.32819 | -7.266 | <0.0001 |
| trans. prob._0.1 x timing_spring+fall | 0.49002 | 0.31139 | 1.574 | 0.116 |
| trans. prob._0.1 x area_20km | -2.02252 | 0.2496 | -8.103 | <0.0001 |
| trans. prob._0.1 x area_40km | -0.66244 | 0.24733 | -2.678 | 0.007 |
| prop. vaccinated x timing_spring | 27.19396 | 0.90888 | 29.92 | <0.0001 |
| prop. vaccinated x timing_spring+fall | -4.81453 | 1.29356 | -3.722 | 0.0002 |
| prop. vaccinated x area_20km | 7.25877 | 0.61762 | 11.753 | <0.0001 |
| prop. vaccinated x area_40km | 2.563 | 0.61626 | 4.159 | <0.0001 |
| timing_spring x area_20km | -2.31549 | 0.26872 | -8.617 | <0.0001 |
| timing_spring+fall x area_20km | 0.65203 | 0.25932 | 2.514 | 0.012 |
| timing_spring x area_40km | -0.3702 | 0.25898 | -1.429 | 0.153 |
| timing_spring+fall x area_40km | 0.02349 | 0.20746 | 0.113 | 0.901 |
| n= observations = 39,600 |  |  |  |  |
| K = number of parameters = 27 |  |  |  |  |
| R<sup>2</sup> = 0.91 |  |  |  |  |

## II. SUPPLEMENTAL METHODS

We describe our modeling approach here using the updated Overview, Design Concepts, and Details protocol for individual-based models (Grimm et al., 2006, 2010).

### A. Overview

#### i. Purpose

Our main objectives were to 1) investigate the effect of variation of host movement distributions on rabies epidemiology in a wild carnivore population, 2) explore the effect of wildlife host movement on the probability that rabies virus will breach an oral rabies vaccination (ORV) zone, and 3) identify the relative effectiveness of feasible components of ORV strategies while accounting for realistic ecological processes in host demography and rabies epidemiology. The components of ORV strategies that we investigated included: seasonal timing of ORV application, width of the vaccination zone, and fraction of wild animals immunized. We first conducted a sensitivity analysis by varying weekly host movement distributions and disease transmission probabilities to explore the effect of variable movement on epidemiological processes. We then examined the effects of host movement variation on ORV effectiveness by comparing epidemiological outputs from two realistic host movement distributions with different variances but similar magnitudes of host movement, and using a full factorial design on the three ORV strategy components.

#### ii. Entities, state variables, and scales

We modeled raccoon individuals as entities with the following attributes: age, sex, group, and individual raccoon identification (ID), natal dispersal age and status, litter size, reproductive status, longevity, home range centroid (x, y coordinates), and grid cell ID. Epidemiological state variables included susceptible, exposed, infectious, and recovered disease classes. Each grid cell represented 1km^2^ and the total area of the modeled landscape ranged between 820km^2^ and 1620 km^2^ depending on the width of the vaccination zone (20, 40, and 60 km, Figs. 1A & 2). Each cell had a carrying capacity of 15 individuals, and was based on published raccoon densities from suburban habitats (Urban 1970; Sonenshine and Winslow 1972; Moore and Kennedy 1985, Table 1). Each raccoon family group had a maximum of 10 individuals. Individuals in the same family group were assigned the same grid cell ID and home range centroid point from which movement originated. Sex and natural longevity were assigned at birth, while all other states changed depending on age, sex, population density, or exposure to an infectious individual.

### B. Process overview and scheduling

Models were updated on a weekly time step. We ran ten-year vaccination simulations and eight-year sensitivity simulations. In each simulation, we initiated disease by seeding one infectious individual into the seeding zone during week 10, following a one-year period for demographic burn-in (i.e., seeding rabies virus in week 63). During the post-burn-in simulation, the order and processes per time step were:

- Update ages and reproductive clocks
- Weekly movement and disease state transitions: randomly assign each individual a weekly home range radius relative to their home range centroid. Transition susceptible individuals to exposed class based on within or between group transmission probabilities given the infectious individuals in their home range at week *t* (see Equation 1 below). Update disease states of exposed and infectious individuals, if applicable.
- Vaccination: randomly select a fixed proportion of the population within the ORV zone; update their disease state to recovered.
- Natural mortality: remove individuals that reach longevity age.
- Natal dispersal: change the home range centroid and grid cell ID (if applicable) of dispersal-age males that group together and dispersal-age males that become solitary (procedure described below). Update natal dispersal status to complete.
- Social dynamics: for family groups that become too large (i.e., greater than 10 individuals), disperse half to another home range centroid and grid cell ID (if applicable) using the same procedure as natal dispersal.
- Density-dependent mortality: for grid cells that exceed the fixed carrying capacity, remove youngest individuals.
- Conception: identify new conceptions, initialize gestation clocks, choose litter size, turn-off postnatal clocks.
- Births: when gestation is complete, assign attributes to each new litter (size determined at random from normal distribution, Table 1). Assign sex in 1:1 ratio, assign the grid cell ID, group ID and home range centroid as mother; assign other individual-level factors.

### C. Design concepts

#### i. Basic principles

We explicitly modeled raccoon population dynamics and RABV transmission on the landscape in two separate analyses. In a first set of simulations (sensitivity analysis), we investigated the effect of variable movement on disease processes by varying weekly host movement distributions and disease transmission probabilities in the absence of vaccination. In the second set of simulations (vaccination analysis), we explored how three components of ORV strategies— proportion of the population vaccinated, timing of ORV, and width of the ORV zone (Fig. 2)—affected vaccination effectiveness. We also examined how data-informed weekly raccoon movement patterns affected ORV zone breach probability because raccoons exhibit variation in home range sizes across different habitats (Beasley & Rhodes, 2010; Šálek, Drahníková, & Tkadlec, 2015).

#### ii. Emergence

Natal dispersal and overcrowding dispersal distances were defined by a random distribution with set parameters, but the realized dispersal distance emerged from both the random distribution and the spatial distribution of current population density. Model algorithms forced individuals to move twice as far as the randomly drawn dispersal distance if the grid into which the individual was moving was already at the grid-level carrying capacity. New distances were considered until either an individual dispersed off the grid permanently or found a grid cell with space.

#### iii. Sensing and Interaction

Raccoons were capable of sensing social group dynamics, within group density, and grid-level density. Susceptible and infectious individuals interacted to determine disease transmission. We did not incorporate explicit interaction of wildlife managers and raccoons.

#### iv. Stochasticity

Longevity, weekly movement distance, dispersal distances, dispersal age (males only), litter size, and disease incubation period were random distributions.

#### v. Collectives

Raccoons were born into family groups. Males remained within family groups until natal dispersal. Females remained in family groups until group size exceeded the group carrying capacity, if applicable.

#### vi. Observation

For the sensitivity simulations, total cases by sex and age, total population size, and the linear distance the pathogen moved were recorded on a weekly timescale. For the vaccination simulations, the same observations were recorded at a quarterly time step, as well as immunity by age class and the total cases within the vaccination zone.

### D. Initialization

The simulated landscape was initialized with raccoons equal to the carrying capacity of each cell. Sex was randomly assigned in a 1:1 ratio. Longevity and age were assigned randomly. We initialized group structure by grouping females that were the minimum reproductive age or older with individuals that were younger than the minimum dispersal age of males. Individual and group IDs were assigned at random, ensuring that there was at least one reproductive female per group and that the total number per group did not exceed the group carrying capacity. Home range centroid coordinates for family groups and males aged 1.5 years and older were randomly chosen within that grid cell. All individuals were assigned initially to the susceptible disease class. Prior to disease introduction, 10% of individuals of sufficient age were randomly chosen and transitioned to the recovered disease class to model natural rabies resistance observed in raccoon populations (Slate et al., 2009). Populations were allowed to undergo demographic dynamics for one year, after which ∼15 infectious individuals were introduced into the seeding zone on week 63 of the simulation.

### E. Input data

Input parameters are described in Table S1. We also input landscapes describing the grid cell IDs, locations, and zonal information. The landscape contained 4 distinct zones (Fig. 1): seeding (1 × 20 km), spreading (10 × 20 km), vaccination zone (20-60 × 20 km), and breach (10 × 20 km).

### F. Submodels

#### i. Natural mortality

Natural mortality was modeled as a random variable (Table S1). Longevity was assigned to individuals at birth, and was drawn from an exponential distribution (values rounded to nearest week; mean = 3 years; Table S1). Individuals were permanently removed at the age of longevity. Individuals were also subject to density-dependent mortality, as well as disease-induced mortality if they became infected and infectious.

#### ii. Social structure

We modeled females and young in family groups. We assumed that the maximum group size was 10 (maximum of 2 adult females and 4 kits/dam). Conception probability varied by age and gestation period was 9 weeks (Table S1). Litter size was drawn from a normal distribution (values rounded to integers; mean = 4 kits; Table S1). Longevity was drawn from an exponential distribution (values rounded to nearest week; mean = 3 years; Table S1). Dispersal-age males were either in male-only dyads or were solitary following a second dispersal event at 1.5 years of age. Raccoons exhibit variable degrees of sociality, with field and genetic studies suggesting that females are often philopatric, and daughters may associate with mothers and offspring into adulthood (Cullingham et al., 2008; Dharmarajan et al., 2009; S. Gehrt & Fritzell, 1998b). Natal dispersal is male-biased, and males may associate in relatively long-lasting, non-familial dyads followed by separation to become independent as they mature (S. D. Gehrt, Gergits, & Fritzell, 2008; S. Gehrt & Fritzell, 1998a; Hirsch, Prange, Hauver, & Gehrt, 2013).

#### iii. Dispersal and weekly movement

We modeled weekly movement as a random variable (gamma distribution, Table 1). Each susceptible individual was assigned a weekly movement distance radius *r* relative to the home range centroid of that group, dyad, or solitary individual. Home range area with radius *r* defined the contact zone for infectious and susceptible individuals, and varied from week to week for susceptible individuals. We assumed that individuals would explore the entirety of the time-varying home range area, and that all contact opportunities within that range area were equally likely.

Natal dispersal was male-biased (Gehrt & Fritzell, 1998). Dispersal distance and natal dispersal age were drawn from a Weibull distribution (mean = 13.6km; Fig. S6, Table S1) and a Poisson distribution (mean = 10 months; Table S1), respectively. Males initially dispersed as male only dyads, and dispersed again at 1.5 years, becoming independent with separate home range centroids. For dispersal due to group overcrowding (>10 individuals, Table S1), groups split into two, with one group dispersing randomly. we modeled two types: post-natal dispersal by males, and relocation by both males and females due to family group overcrowding (Fig. 2, Table 1). Movement into cells at grid-level carrying capacity was not allowed. Dispersal events occurred as follows. For each 45-degree angle from the home range centroid, movement distances were chosen at random from the specified random distribution of weekly movement distances. New potential x, y coordinates were calculated as x = distance × cos(angle) + current × coordinate, y = distance × sin (angle) + current y coordinate. We chose 45-degree angles to systematically search all surrounding neighbor grid cells at the randomly drawn dispersal distance (Queen’s case contiguity). Dispersal occurred if the target locations were located within the simulated landscape and within a grid with fewer raccoons than the carrying capacity. When more than one location satisfied the criteria, the new location was chosen at random among those that were considered valid. If no locations were valid, the distance values were doubled, and the process repeated until a valid set of coordinates were found (or the individual moved off the landscape. We thus assumed that dispersing individuals would move farther and farther out in a crowded landscape, reflecting(Bowler, Benton, Bowler, & Benton, 2015) evidence that high density populations can decrease the immigration success of some mammal species Individuals that traveled off the landscape were lost permanently and did not return to the landscape.

#### iv. Disease transmission

We modeled epidemiological dynamics of rabies with four disease states: susceptible, exposed but not infectious, infectious, and recovered (Figs. 1B & S1A), as detailed in the methods of the main text. Incubation period of exposed individuals was drawn from a Poisson distribution with a mean of 4 weeks (Tinline *et al*. 2002; Rees *et al*. 2008, Table 1). Disease-induced mortality for infectious individuals was 100% and death from disease occurred one week after individuals transitioned from the exposed class into the infectious disease class (Hanlon et al., 2007). Recovery rate of exposed individuals was 10% to capture variation in natural resistance to rabies observed in raccoons (Slate et al., 2014). Susceptible individuals that were vaccinated entered the recovered disease class with a two-week lag following the vaccination activity.

#### v. Vaccination

Vaccination comprised three components: coverage or proportion of animals immunized within the ORV zone, timing and frequency of vaccination, and ORV zone width (Fig. 2). We examined three zone widths: 20 km, 40 km, and 60 km, with larger widths implemented as additional grid cells in the ORV zone of the landscape. We modeled vaccination coverage by randomly selecting a fixed proportion of the population within the ORV zone, and converting susceptible individuals within this subset to the recovered disease class with a delay of 2 weeks to account for the development of immunity in vaccinated individuals. Raccoons under 17 weeks are still nursing (Montgomery, 1969); only individuals older than 17 weeks could be vaccinated because delivery was assumed to be through oral vaccine baiting. Vaccination coverage ranged from 0 - 100% where 0% was the no vaccination control (Figs. 2 & S2). Raccoon ORV was timed to occur during fall, spring, or both fall and spring to explore different strategies for timing and frequency because seasonality in host or pathogen ecology can drive disease dynamics such that intervention timing and frequency may influence intervention effectiveness (Altizer et al., 2006). Vaccination was assumed to be 100% effective for susceptible and exposed individuals and imparted lifetime immunity. Vaccination was assumed ineffective on infectious individuals.

#### vi. Density-dependent mortality

In order to limit population abundance to our given within-grid carrying capacity, we implemented density-dependent mortality. When grid cells exceeded the carrying capacity (15 individuals per km^2^), individuals within the grid were randomly chosen and removed from the simulation, with younger individuals taken first to mimic observed differences in juvenile and adult survivorship (G. Gehrt & Fritzell, 1999). We assumed no time lag between when grids exceeded the within-grid carrying capacity and the resulting effects on mortality of individuals within that grid.

#### vii. Conceptions and births

We modeled reproduction as a single birth pulse beginning in April and extending to mid-May for a 6 week breeding period (George & Stitt, 1951; Rees et al., 2008; Stuewer, 1943). Conception probability varied by age and gestation period was 9 weeks (Table S1), thus mating occurred from mid-February to early March. The probability of conception varied with age such that females > 1.5 years of age were more likely to conceive relative to females < 1.5 years of age, implemented using scaling parameters on conception probability (Table 1). Litter size was drawn from a normal distribution (values rounded to integers; mean = 4 kits; Table S1). Litter size was a drawn from a normal distribution with a mean of 4 kits and variance = 1 (Sanderson and Hubert 1981; Fritzell *et al*. 1985; Ritke 1990; Table 1).

### G. Movement data collection and fitting

We fit weekly distances to a gamma distribution and estimated the shape and scale parameters (shape= 4.1 and scale= 0.2) using maximum likelihood methods. The mean and median of the gamma distribution fit to these data was 0.82 km and 0.75 km, respectively, and the variance was the probability that any individual moved more than 3 km was 0.00001. We used a second gamma distribution with a higher mean, median, and variance (shape = 2, scale = 0.5, mean = 1 km, median = 0.84) to explore the effect of a more variable host movement distribution on vaccination effectiveness; the probability any individual moved more than 3 km was approximately 0.018—substantially higher than the less variable movement distribution.

We estimated similar shape and scale parameters from an independent study of GPS-collared raccoons in a suburban ORV area in Tennessee over the same months during 2013-2014 (Berentsen et al., 2017; data not shown). Data collection and fitting procedures are further detailed in the methods in the main text.

### H. Simulations

We conducted outbreak simulations using multiple levels of three parameters: 1) shape and 2) scale parameters of the weekly host movement gamma distribution, and 3) between-group transmission probability. We focused on weekly host movement distributions rather than host dispersal movement distributions because results of a preliminary sensitivity analysis (data not shown) indicated that disease outcomes were not sensitive to variability in the natal dispersal variable but were sensitive to variability in the weekly movement variable.

To further investigate the relative effects of magnitude (median) and variability of weekly movement gamma distributions on breach probability, we modeled vaccination using four gamma distributions. Two gamma distributions had similar medians (median = 0.8 km) but differed in variance (variance = 0.49, Gamma(shape=1.96, scale=0.5); variance = 0.17, Gamma(shape=4.3, scale=0.2). Two gamma distributions had similar variance (variance = 0.3) but differed in median (median = 0.75km, Gamma(shape=2.2, scale=0.4; median = 0.87 km, Gamma(shape=2.8, scale=0.35). Vaccination zone area (40km), ORV application time and frequency (fall only), and transmission probability (0.05) were held constant. We ran 43 unique parameter sets, with 100 ten-year replicate simulation per parameter set, for a total of 44,000 simulations. We defined vaccination effectiveness as 1 – *v*, where v is the minimum vaccination coverage required to reduce breach probability to zero.

#### R_0_ Simulation

We calculated R_0_ (the average number of infectious cases caused by one infectious individual in a completely susceptible population) using numerical simulation for one set of conditions that implemented the data-informed gamma distribution (Gamma(shape = 4.1, scale = 0.2)) for weekly home range movement and a transmission probability of 0.05. We conducted 1000 two-year replicate simulations where we tracked the number of transmissions caused by a single index case, and took the mean number of index case transmissions as R_0_.

##### I. Model outputs and statistical analysis

See methods in main text for description of the main model outputs and statistical analyses.

## III. SUPPLEMENTAL RESULTS

### A. Additional vaccination simulation

Increases in the variance of the weekly movement gamma distribution led to a 42.9% decrease in vaccination effectiveness, when median was held constant between movement distributions. Increases in median led to a 16.7% decrease in effectiveness.

